# Organism-wide single-cell transcriptomics of long-lived *C. elegans daf*-2^-/-^ mutants reveals tissue-specific reprogramming of gene expression networks

**DOI:** 10.1101/509992

**Authors:** Jessica L. Preston, Nicholas Stiffler, Maggie Weitzman

## Abstract

A critical requirement for a systems-level understanding of complex biological processes such as aging is the ability to directly characterize interactions between cells and tissues within a multicellular organism. *C. elegans* nematodes harboring mutations in the insulin-like receptor *daf-2* exhibit dramatically-increased lifespans. To identify tissue-specific biochemical mechanisms regulating aging plasticity, we single-cell sequenced 3’-mRNA libraries generated from seven populations of whole day-one adult wild-type and *daf-2*^*-/-*^ worms using the 10x ChromiumV1™platform. The age-synchronized samples were bioinformatically merged into a single aligned dataset containing 40,000 age-synchronized wild-type and *daf-2*^*-/-*^ cellular transcriptomes partitioned into 101 clusters, using unsupervised machine-learning algorithms to identify common cell types. Here we describe the basic features of the adult *C. elegans* single-cell transcriptome and summarize functional alterations observed in the gene expression profiles of long-lived *daf-2*^*-/-*^ worms. Comprehensive methods and datasets are provided. This is the first study to directly quantify cell-specific differential gene expression between two age-synchronized, genetically-distinct populations of multicellular organisms. This novel approach answers fundamental questions regarding tissue-specific regulation of gene expression and helps to establish a foundation for a comprehensive *C. elegans* single-cell gene expression atlas.

## Introduction

Throughout the development and aging of a multicellular organism, genomic DNA is transcribed into a highly heterogeneous set of individual cellular transcriptomes with a wide range of biological functions. The various cell types communicate through complex chemical and electrical signaling networks in order to fine-tune the physiological response of each individual cell to its unique environmental context. These interconnected non-autonomous cell signaling pathways are tightly-regulated to specifically tailor the metabolic, reproductive, and stress-induced transcriptional response of each individual cell to its current spatiotemporal niche.

Longevity studies in model organisms have demonstrated that biological aging rates are highly plastic and depend on genetic and environmental factors (Herndon, Schmeissner et al. 2002, Artal-Sanz and Tavernarakis 2008, Blagosklonny, Campisi et al. 2010, Kenyon 2010, Kenyon 2010, Lopez-Otin, Blasco et al. 2013, Wu, Liu et al. 2013, Wu, Liu et al. 2013, Zimmerman, Hinkson et al. 2015). Despite the significant progress made towards understanding the basic aging process, the specific biochemical mechanisms regulating aging plasticity and senescence remain largely uncharacterized. Longevity rates are directly and indirectly influenced by numerous signaling networks controlling fundamental physiological processes such as metabolism, reproduction, and immune response. (Lund, Tedesco et al. 2002, Viswanathan, Kim et al. 2005, Budovskaya, Wu et al. 2008, David, Ollikainen et al. 2010, Shin, Lee et al. 2011, Youngman, Rogers et al. 2011, Back, Braeckman et al. 2012, Hou and Taubert 2012, Yashin, Arbeev et al. 2013, Pan, Li et al. 2016). At the systems-level, the global integration of every metabolic decision of every cell ultimately determines the duration of an organism’s lifetime. In order to predict and control the aging rate of an organism, the fundamental regulatory pathways underlying lifespan determination must be identified.

The nematode *C. elegans* is a powerful and well-established model organism for investigating embryonic development, reproduction, and aging. The lifespan of the nematode is short and highly-plastic, and the entire developmental cell lineage and the neuronal connectome of the worm are mapped and stereotypic (White, Horvitz et al. 1982, Sulston, Schierenberg et al. 1983, Sulston 2003, Towlson, Vertes et al. 2013, Arnatkeviciute, Fulcher et al. 2018). *C. elegans* strains harboring temperature-sensitive mutations in the insulin-like receptor *daf-2* have dramatically-extended lifespans due to increased activation of the stress-induced transcription factor *daf-16* (FoxO) (Kenyon, Chang et al. 1993, Murphy, McCarroll et al. 2003, Halaschek-Wiener, Khattra et al. 2005, Patel, Garza-Garcia et al. 2008, David, Ollikainen et al. 2010, Henis-Korenblit, Zhang et al. 2010, Kaletsky and Murphy 2010). The biochemical mechanisms underlying the lifespan extensions characteristic of *daf-2* hypomorphs are difficult to decipher due to the pleiotropic nature of *daf-2* alleles. Organism-wide knockdown of *daf-2* causes widespread and extensive disruptions in several ubiquitous cell signaling networks (Jia, Chen et al. 2004, Back, Braeckman et al. 2012, Qi, Huang et al. 2012, Wan, Zheng et al. 2013, Ewald, Landis et al. 2015). Cross-talk between diverse cell types induces complex non-autonomous and tissue-specific metabolic responses which are unable to be resolved with standard bulk RNA-Seq methods. Progress in *C. elegans* tissue-specific transcriptomics was previously hindered due to technical challenges arising from the nematode’s tough outer cuticle, as well as the lack of robust antibodies or methods available for sorting of live *C. elegans* cells with flow cytometry. Recent breakthroughs in worm dissociation techniques (Fernandez, Mis et al. 2010, Zhang, Banerjee et al. 2011, Kaletsky, Lakhina et al. 2016) have enabled high-throughput characterization of individual cells from whole adult nematode.

Single-cell RNA-Seq (sc-RNA-Seq) is a powerful tool for resolving the transcriptional heterogeneity of complex tissues down to the level of the individual cell, revealing previously undetectable signals in gene expression data (Buettner, Natarajan et al. 2015, Levine, Simonds et al. 2015, Macosko, Basu et al. 2015, Satija, Farrell et al. 2015, Scialdone, Natarajan et al. 2015, Xu and Su 2015, McKenna, Findlay et al. 2016, Cao, Packer et al. 2017, Saunders, Macosko et al. 2018). The completely-mapped cell lineage of *C. elegans* makes it an ideal model organism for single-cell expression studies. The extensive knowledge acquired from decades of meticulous investigations into the developmental origin, function, position, and anatomical features of each of the worm’s 959 adult somatic cells establishes a solid foundation on which to build a comprehensive atlas of organism-wide transcriptional programming. Here we show that droplet-based 10x ChromiumV1™ single-cell barcoding technology (Zheng, Terry et al. 2017) can be applied to capture transcriptomic data from individual cells of whole adult nematodes. This technology was used to characterize the tissue-specific gene expression profiles of wild-type and long-lived *daf-2*^*-/-*^ mutant *C. elegans* populations, in order to decipher systems-level regulatory mechanisms controlling aging rate determination.

## Results

### I. Whole-worm single-cell 3’-mRNA-sequencing

To achieve a comprehensive and unbiased view of the adult *C. elegans* transcriptome, we employed whole-worm single-cell 10x Genomics^®^ ChromiumV1™3’-mRNA-sequencing combined with unsupervised machine learning-based characterization of cell-specific gene expression patterns (Zheng, Terry et al. 2017, Butler, Hoffman et al. 2018). Suspensions of live dissociated *C. elegans* cells were prepared from age-synchronized populations of whole day-one adult wild-type and *daf-2*^*-/-*^ worms as described, using a 5-micron filter (Kaletsky, Lakhina et al. 2016, Kaletsky, Yao et al. 2018). Live cells were individually-partitioned into aqueous oil-emulsion nanodroplets using 10x ChromiumV1™microfluidics technology. Single cells were chemically lysed while inside the droplets, and mRNA transcripts were captured via polyA-tail hybridization to oligonucleotide-barcoded gel beads. Individual transcripts were tagged with a unique combination of three barcodes to facilitate downstream tracking of the genetic, cellular, and molecular origin of each sequencing read. Three prime mRNA sequencing (3’-mRNA-Seq) of the final barcoded 10x ChromiumV1™single-cell libraries was conducted on an Illumina NextSeq500 ™using 75 basepair (bp) reads. The default 10x CellRanger ™(v1.2.1) software pipeline was used to perform genome alignment, barcode quantification and filtering, duplicate read removal, gene expression normalization using unique molecular identifier (UMI) tags, and cell count determination.

### II. Characterization of wild-type and *daf-2*^*-/-*^ mutant adult *C. elegans* cellular transcriptomes

Single-cell transcriptomic datasets generated from seven independent age-synchronized populations of adult *C. elegans* were bioinformatically aligned and merged into a single sc-RNA-Seq dataset using unsupervised machine-learning algorithms (Waltman and van Eck 2013) implemented with the Seurat (v2.0.1) canonical correlation dataset alignment (CCA) pipeline. After normalization, single-cell transcriptomes were digitally partitioned into distinct groups based on transcriptional similarity, using Seurat (v2.0.1)-implemented unsupervised hierarchical clustering algorithms (Butler, Hoffman et al. 2018). The final aligned dataset contained ∼40,000 age-synchronized wild-type and *daf-2*^*-/-*^ cells divided into 101 statistically-distinct cell type clusters (Figure 1). Each machine-generated cell type cluster is characterized by the expression of a unique group of biomarkers which is stable across replicates, known as its gene expression signature (Table S1). The cell type clusters in the aligned dataset are comprised of a heterogeneous mixture of cells from seven independent populations of age-synchronized wild-type and *daf-2*^*-/-*^ worms (Table S2). While the majority of the cell type clusters contain both wild-type and *daf-2*^*-/-*^ cells, twenty small clusters contain primarily cells of a single genotype (Table S3).

**Figure 1.**
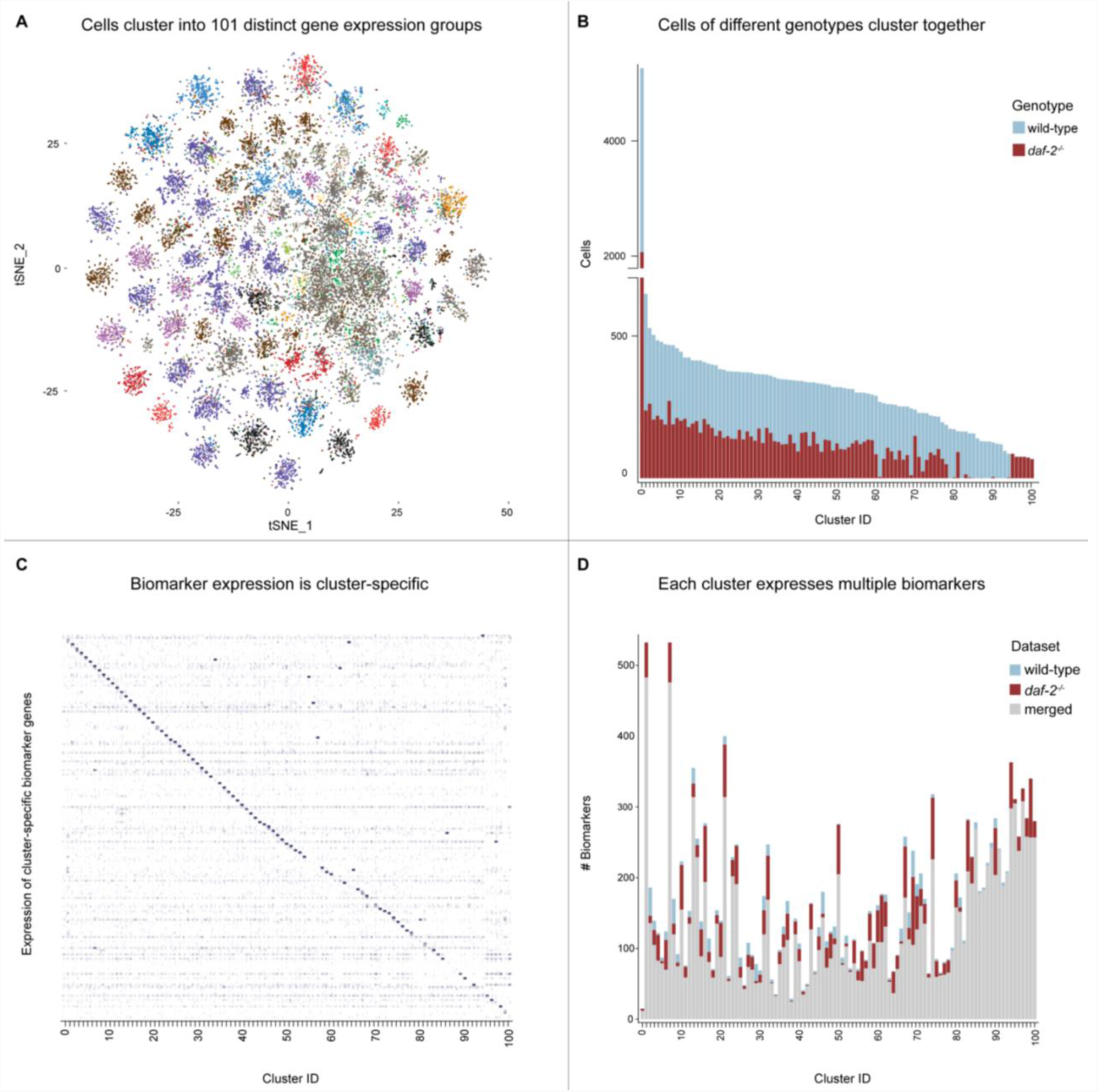
Unsupervised hierarchical clustering of individual cellular transcriptomes from whole day-one adult *C. elegans*. For this study, datasets from seven distinct populations sequenced over four months were combined using unsupervised sample alignment algorithms. After sample normalization, the final dataset contained ∼40,000 cells partitioned into 101 cell type clusters, each with its own unique biomarker signature **A**. A tSNE plot of ∼40,000 single-cell transcriptomes from seven populations of whole adult *C. elegans* day one *daf-2-/-* and *wildtype* adults. **B**. Cell counts present in each cluster. Most of the 101 cell type clusters contain cells of both genotype. **C.** Biomarker specificity. Each cluster is identifiable based on strong expression of unique biomarker genes. The biomarker signatures are highly statistically significant, with adjusted p-values approaching ∼0. Biomarker signatures are highly-specific to their respective cell cluster, often exhibiting undetectable expression levels in all other cell types. **D.** Biomarker counts by cluster. The cluster biomarker profile was determined for each genotype individually (blue and red bars) and for the entire merged dataset (grey bars).

### III. Cell type determination and biomarker discovery

Cells were grouped into cell lineage clusters based on similar gene expression profiles. *C. elegans* single-cell clusters express distinct sets of marker genes that are remarkably consistent and reproducible over time. The biomarker genes identified in this study are stably expressed across biological replicates and highly statistically significant. Several marker genes have adjusted p-values reported as approximately equal to zero by the Seurat(v2.0.1):FindMarkers pipeline. The predicted cell types reported in this study (Figures 2,3; Table S4) were assigned to each cluster using a supervised approach. Cell types were inferred for each expression group by integrating the documented functional and anatomical information of each biomarker gene with its experimentally observed specificity and significance. The biomarker lists generated by the software were analyzed using the WormBase SimpleMine database, which provides high-throughput descriptions of gene function and tissue-specific expression. Several genes were identified which provide anatomical, biochemical, and genetic information about the cell type clusters supported by the literature (Tomaselli, Reichardt et al. 1986, Way and Chalfie 1989, Way, Wang et al. 1991, Gendreau, Moskowitz et al. 1994, Wang and Way 1996, Tavernarakis and Driscoll 1997, Tavernarakis, Shreffler et al. 1997, Duggan, Ma et al. 1998, Koh and Rothman 2001, Jia, Chen et al. 2004, Fukushige, Brodigan et al. 2006, Bacaj, Lu et al. 2008, Smit, Schnabel et al. 2008, Barrios, Ghosh et al. 2012, Frooninckx, Van Rompay et al. 2012, Goodwin, Sasaki et al. 2012, Komuniecki, Harris et al. 2012, Sasidharan, Sumakovic et al. 2012, Jedrusik-Bode 2013, Towlson, Vertes et al. 2013, Ackley 2014, Ortiz, Noble et al. 2014, Peymen, Watteyne et al. 2014, Burdick, Walton et al. 2016, Kaletsky, Yao et al. 2018).

**Figure 2.**
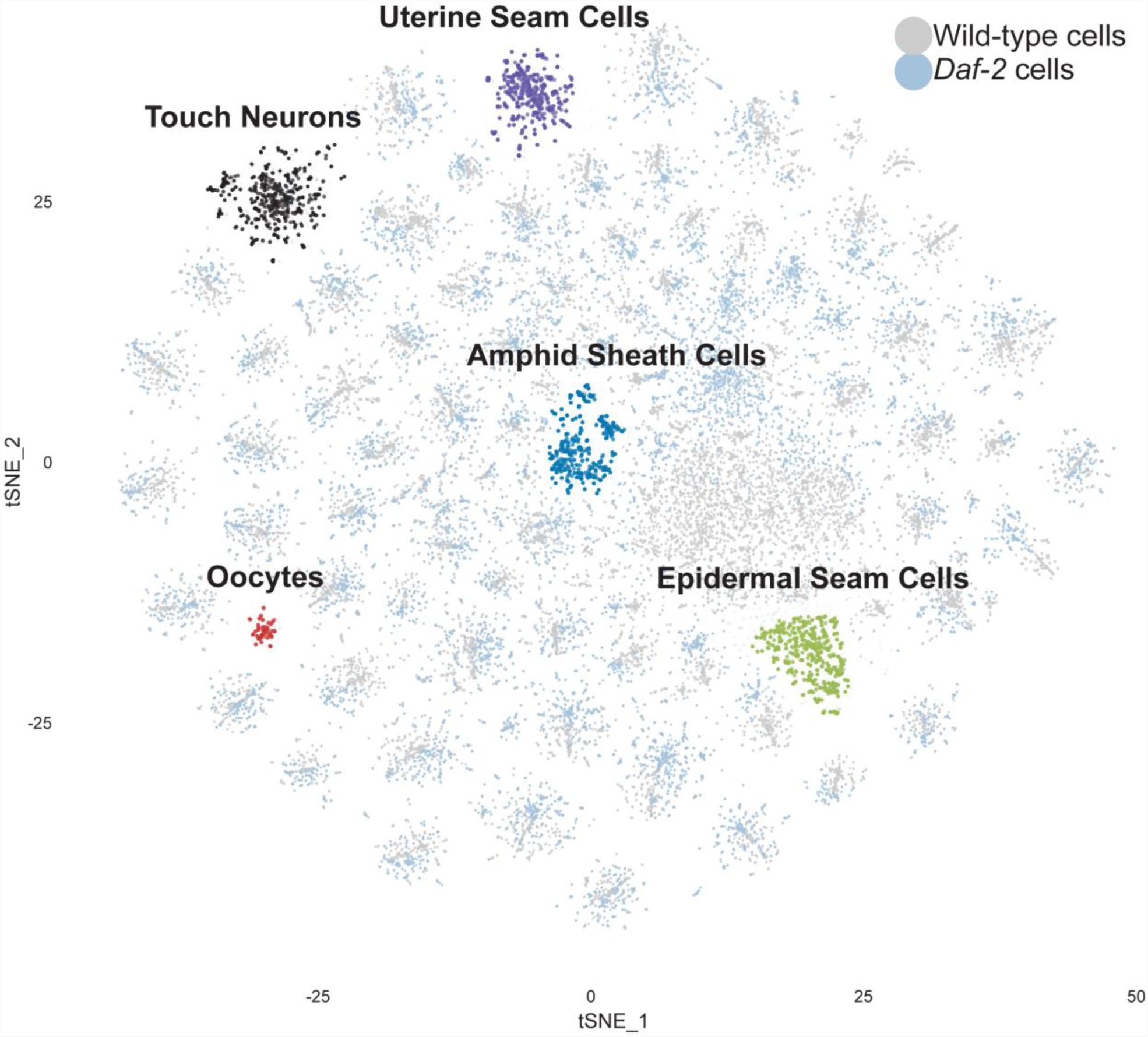
A tSNE plot depicting ∼40,000 single cells from seven independent samples of age-matched wild-type and *daf-2-/-* adults *C. elegans* worms grouped into 101 aligned cell type clusters using unsupervised hierarchical clustering of gene expression profiles. Representative tissue types are labelled with predicted functional identities.

**Figure 3.**
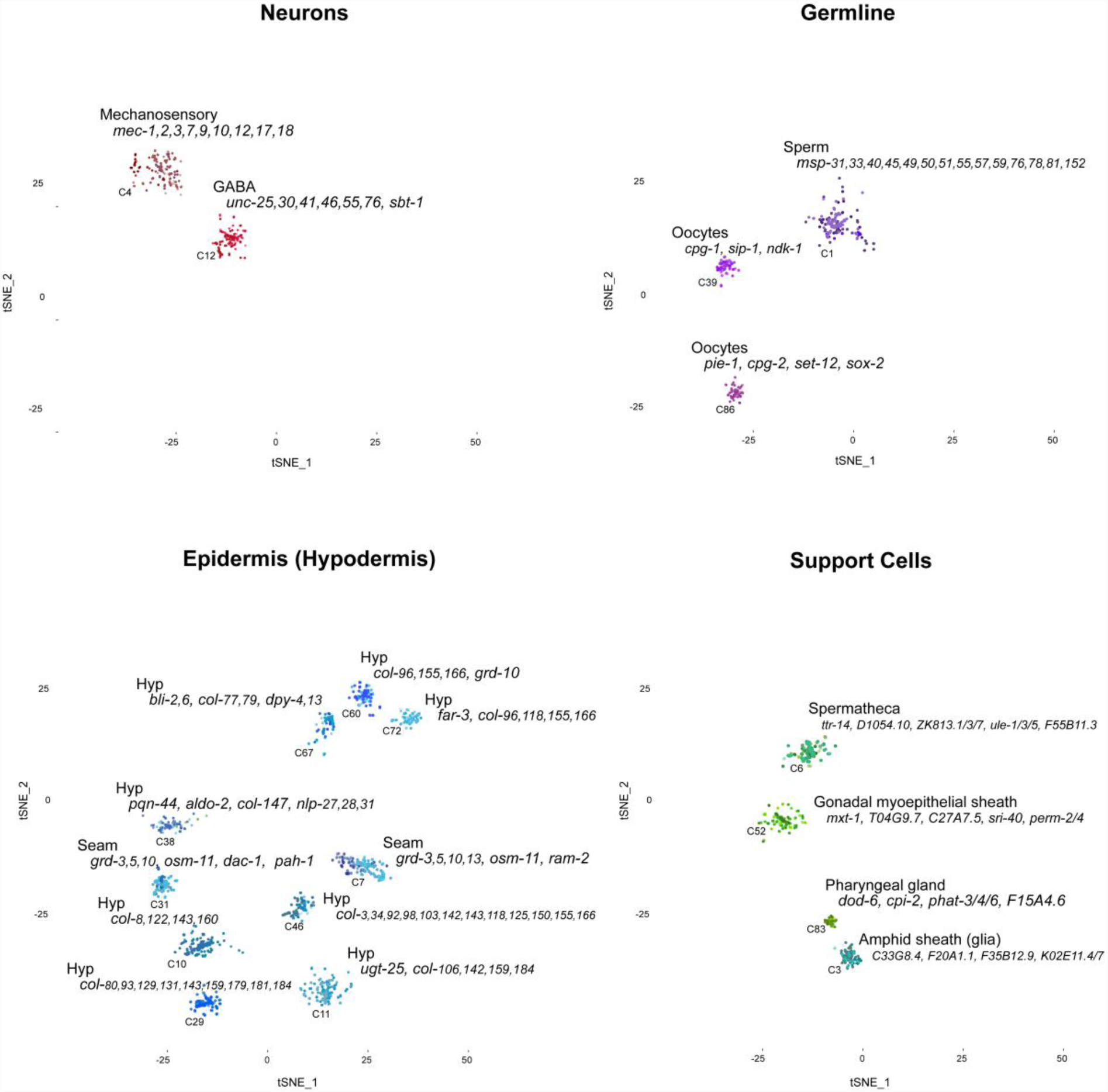
Tissue-specific biomarker profiles in wild-type adult *C. elegans.* Gene expression signatures of various representative tissue-type clusters are illustrated. Many cell clusters express biomarker signatures comprised entirely of novel genes, including amphid sheath (glia) and spermatheca cell clusters.

Several novel tissue-specific biomarker genes were identified which are completely uncharacterized. Interestingly, the expression signature of a large, dense cluster of amphid sheath cells (Cluster #3) is composed almost entirely of genes with unknown functions (Table S5) (Bacaj, Tevlin et al. 2008).

Some *C. elegans* single-cell clusters exhibit co-expression of biomarker genes reported as being expressed in developmentally-related tissues. For example, many predicted sheath cell clusters contain several marker genes expressed in both cephalic sheath cells and spermatheca, which share a common developmental lineage (L’Hernault 2006). In addition, many *C. elegans* single-cells cluster based on the co-expression of multiple marker genes that are consistently grouped together in different cell types. For example, *lec-8, dct-16, ftn-2, C17F4.7, Y38F1A.6, F48D6.4, and Y119D3B.21* are often co-expressed in the intestine and pharynx. These genes may function in conserved terminal differentiation programs that operate throughout development in distinct tissue types.

### IV. Tissue-specific transcriptional responses to global *daf-2* knockdown

Automated quantifications of cell-specific differential gene expression were made possible with the Seurat (v2.0.1) canonical correlation dataset alignment (CCA) procedure. The CCA employs machine-learning algorithms to identify and combine common cell types across the age-synchronized datasets (Butler, Hoffman et al. 2018). After global scaling and normalization of gene-expression data across all cell types and samples, cell-specific differential gene expression patterns can be directly calculated using a single, integrated high-throughput pipeline. The dataset alignment procedure directly compares the gene expression profiles of specific cell types present in wild-type and *daf-2*^*-/-*^ worms. Systems-level high-throughput quantification of cell-specific differential gene expression based on genotype was performed on the 81 mixed-genotype clusters using the Seurat(v2.0.1):FindMarkers function, which calculates differential expression between any two groups of cells. Genes with significantly altered expression between wild-type and *daf-2*^*-/-*^ transcriptomes were calculated for each cluster individually using a threshold of padj < 0.10 to define significance (Supplemental File 2).

The global downregulation of *daf-2* gene function led to widespread and complex effects on tissue-specific gene expression throughout the organism (Figure 4). Collagens are well-represented in the tissue-specific transcriptional responses to global *daf-2* knockout, exhibiting extremely strong expression dynamics. Significantly altered expression was detected in functional gene networks involved in extracellular matrix remodeling (collagens), innate immunity, stress response, protein aggregation, chromatin remodeling, lipid signaling, DNA repair, membrane dynamics, and reproduction (Table S6) (Spieth, Denison et al. 1985, Sherman-Baust, Weeraratna et al. 2003, Haskins, Russell et al. 2008, Schnoor, Cullen et al. 2008, Schulenburg, Hoeppner et al. 2008, Passos, Nelson et al. 2010, Li, Patterson et al. 2012, Studencka, Konzer et al. 2012, Studencka, Wesolowski et al. 2012, Chen, Cescon et al. 2013, Sen, Kundu et al. 2013, Stroehlein, Young et al. 2016).

**Figure 4.**
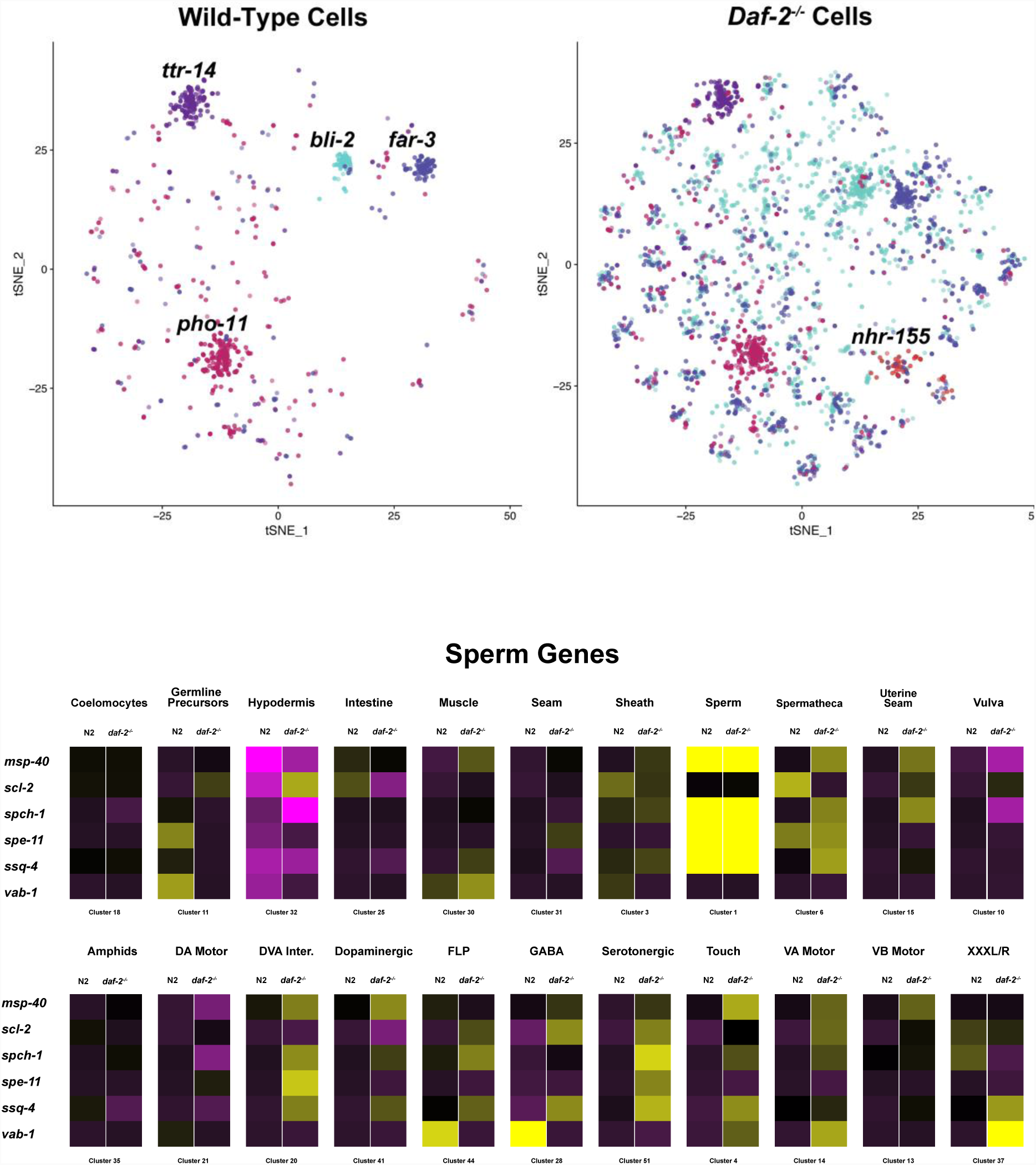
Inducible loss of daf-2 modifies tissue-specific gene expression patterns in adult C. elegans. The global downregulation of *daf-2* gene function led to widespread and complex effects on tissue-specific gene expression throughout the organism.

The unique transcriptional response of each cluster due to the knockdown of *daf-2* was compared with all other clusters to differentiate between the tissue-specific and global effects of *daf-2* knockdown. In addition to the unique tissue-specific responses to daf-2 knockout, there is also a global (organism-wide) gene expression response to global knockdown of *daf-2*. Twenty-seven genes were identified that are significantly differentially expressed in every cluster (Table S7). The global *daf-2*-response genes include several collagens that are highly upregulated in *daf-2* tissues. Unexpectedly, five vitellogenin (yolk protein) genes were identified that are ubiquitously upregulated in all wild-type clusters.

The epidermal seam cells (Koh and Rothman 2001) displayed the strongest response to the loss of *daf-2* in terms of the number of significantly upregulated age-related genes. Dozens of genes are highly expressed in the lateral seam cells of *daf-2*^*-/-*^ animals, including several hormone-induced transcription factors (*nhr-* gene family), groundhog-like genes (*grd-* gene family), and a large network of genes involved in endosomal packaging and distribution of neuromodulators (Table S8). These result suggest that seam cells facilitate non-cell autonomous signaling via endosomal diffusion of neuromodulators (Kaletsky, Yao et al. 2018), possibly through a sterol-induced mechanism of extracellular matrix remodeling which facilitates increased vesicle trafficking.

A wild-type specific cluster of pharyngeal gland cells (Cluster #83) is distinguishable based on its upregulation of the *daf-16*-target gene *dod-6* (“downstream-of-*daf-16*”-6), which is generally upregulated in *daf-2*^*-/-*^ tissues (Murphy, McCarroll et al. 2003). Also unexpected was the discovery that several sperm-specific genes were strongly upregulated in somatic tissues of *daf-2*^*-/-*^ worms, supporting the possibility of a soma-to-germline transformation in the transcriptome of *daf-2*^*-/-*^ worms (Table S9, Figure 4) (Curran, Wu et al. 2009).

## Discussion

Sequencing libraries were generated from heterogeneous mixtures of live dissociated worm cells, directly after filtering with a 5 micron filter. The original intention was to capture only neurons with the 5 micron filter in order to concentrate focus and increase chances of obtaining interpretable data. Interestingly, this method appears to preserve the cellular heterogeneity of the worm, as every tissue type is represented in the data. This non-targeted approach is less biased than antibody-based tissue-specific transcriptomics methods, which rely heavily on *a priori* assumptions classifying cell types based on the absolute (discrete) expression of a single, sometimes poorly-characterized, antigen.

By quantifying the expression levels of the highly-expressed genes in *C. elegans* single cells and performing unsupervised hierarchical clustering, we found that the cells grouped reliably into clusters based on transcriptome similarity. This method was highly reproducible over time and across replicates. Library preparation was performed in two rounds several months apart using worms with slight differences in worm age (∼6 hours), and qualitatively indistinguishable cell clusters were observed in the data.

The number of single-cell clusters identified in a sample using unsupervised clustering was highly dependent on the number of cells present in the original sample, with larger cell numbers leading to more clusters. Straightforward cell type identification was hindered by several obstacles, including the current lack of well-characterized cell-specific promoters in the *C. elegans* genome, the immense complexity of single-cell datasets generated from whole worms, the suboptimal genomic coverage of the ChromiumV1™libraries, and the subjective and time-consuming process of synthesizing information from published datasets. Most *C. elegans* single-cell clusters were easily identifiable based on clear expression of a distinct set of marker genes, allowing identification of the anatomical region or biochemical function of each cell type. However, many clusters contained marker genes with an unclear relationship, annotated as being expressed in several distinct cell and tissue types. The future generation of larger and more comprehensive single-cell datasets by the broader scientific community should help to streamline this process.

The vast scale, complexity, and richness of whole-worm single-cell datasets is breathtaking, yet overwhelming. In order to expedite the unrestricted release of our methods and datasets as a resource to the broader scientific community, the scope of the results reported here were condensed to highlight just the main features of the functional alterations observed in the gene expression profiles of long-lived *daf-2*^*-/-*^ worms.

Most cell type clusters contain both wild-type and *daf-2*^*-/-*^ cells, but twenty of the smallest clusters contain cells of a single genotype. The presence of genetically homogenous cell clusters in the aligned and dimensionally-reduced subspace implies the existence of specific cell types with unique functional differences between the wild-type and *daf-2*^*-/-*^ animals. Despite the genotype-specific condition of these clusters, their transcriptional profiles were similar enough to be recognized and incorporated into the aligned subspace by the unsupervised software.

It is unclear whether cellular function or developmental origin has more impact on cell clustering based on gene expression. Many cell clusters have expression profiles dominated by large interconnected gene networks that are specifically co-expressed in tissues with shared developmental lineages. However, the complex relationship between cell function, gene expression and developmental cell lineages in *C. elegans* requires further investigation.

This is the first study to directly quantify cell-specific differential gene expression between two age-synchronized, genetically-distinct populations of multicellular organisms. This novel approach answers fundamental questions regarding the tissue-specific regulation of gene expression during aging, and establishes the foundation for a comprehensive *C. elegans* single-cell gene expression atlas.

## Methods

### Worm cultivation and strains

*C. elegans* strains were maintained on NHGM plates on *E. coli* OP50 at 20^°^C. Prior to cell dissociation, L1 larvae were age-synchronized using hypochlorite treatment and plated at 15°C to avoid dauer formation in the temperature-sensitive *daf-2*^*-/-*^ mutant strains. Worms were shifted to the *daf-2*^*-/-*^-restrictive temperature of 25°C at L4 stage and grown to day one of adulthood. Strains were provided by the *Caenorhabditis* Genetics Center. OH441: otIs45 [*unc-119*::GFP], CB1370: *daf-2*(e1370), DR1564: *daf-2*(m41), GR1309: *daf-16(mu86); daf-2(e1370).* N2 (Bristol) was used as the wild-type strain.

### Dissociation of adult *C. elegans* into single cells

The 10x Genomics^®^ GemCode™v1.0) scRNA-seq protocol requires live suspensions of single cells as input. Therefore, it is critically important to confirm the viability, purity, and complexity of each sample prior to sequencing in order to obtain viable data. To avoid possible downstream interference with the transcriptional enzymes used during library preparation, all cell dissociation reactions were performed in the absence of the transcriptional inhibitor actinomycin D.

Synchronized day-one adult worms were dissociated into single cells as described previously (Zhang, Banerjee et al. 2011, Kaletsky, Lakhina et al. 2016) with the following modifications. Four large plates of age-synchronized adult worms per sample were washed with 8 µL of ddH2O into a 10 mL centrifuge tube and pelleted by centrifugation at 18,000xg for 1 min. A ∼250 µL pellet of worms was transferred to a 1.6 mL microcentrifuge tube, washed 5X with M9 and then incubated in a lysis buffer (200 mM DTT, 0.25% SDS, 3% sucrose, 20 nM HEPES pH8) for ∼6 min at 20°C. The reaction was closely monitored under a light microscope by regular inspection of 5 µL aliquots and stopped once cuticles were mostly denatured but worms were still intact. Worms were washed 5X with M9 and resuspended in 20 mg/mL pronase enzyme from *Streptomyces griseus* (Millipore Sigma), a non-specific protease. Worms were digested with pronase for 15-20 min at 20°C and mechanically disrupted by rapid pipetting 100X with a regular-bore micropipette tip every 2-3 min, rotating between samples. The dissociation reaction was regularly monitored and stopped with 250 µL of ice-cold 2% fetal bovine serum (FBS) in PBS^-/-^ once cells were visible and the majority of worm heads were dissociated. The worm suspension was pulled into a 1 mL syringe through a 27-gauge needle and then passed through a 5 µm Supor membrane syringe filter (Pall) into collection tubes on ice. Filtered cells were washed twice with 0.04% bovine serum albumin (BSA) in PBS and resuspended in 50 µL of ice-cold BSA/PBS.

### Cell quantification and viability assays

To verify that live neurons could be isolated using this approach, Fluorescence-Activated Cell Sorting (FACS) was performed on dissociated cells from adult *C. elegans* expressing GFP under the control of the *unc-119* promoter (OH441: otIs45 [P*unc-119::GFP*]) (Figure S1). Cells expressing *unc-119* mRNA were isolated based on high GFP protein expression (GFP^high^) using a Sony SH800S flow cytometer with a 70 µm microfluidic sorting chip. DAPI (4’,6-diamidino-2-phenylindole, 1:10^4^) counterstaining was used as a viability marker. Age-synchronized wild-type worms were used as a negative control to set FACS gates. Putative live neurons (DAPI^neg^ / GFP^high^ events) were collected on ice, observed under a fluorescent microscope, and quantified. FACS yields of live C. *elegans* neurons were ∼3% of total live cells, which is within range of previously published values (Fernandez, Mis et al. 2010).

Live *C. elegans* neurons isolated with FACS were assayed immediately following sorting with a Bio-Rad TC20™ automatic cell counter. The viability of the FACS-isolated cells was found to be dramatically lower than non-sorted controls, indicating that most live GFP-expressing neurons were killed during the sorting process. Unlike sci-RNA-seq libraries, which can be prepared directly from non-living cells (Cao, Packer et al. 2017), 10x ChromiumV1 sc-RNA-Seq libraries must be generated using live cells with intact plasma membranes. Because live single cells are required as input for the 10x Genomics^®^ sc-RNA-seq protocol, targeted isolation of *C. elegans* cell types with FACS prior to library generation was subsequently avoided, and sequencing libraries were prepared directly from unsorted whole-worm cell suspensions immediately following 5 µM filtering and washing. Cells were pelleted by centrifugation and resuspended in 50 µL 0.04% BSA in PBS.

Filtered cell suspensions were observed using a Keyence BZ-X800 inverted fluorescence microscope (Figure S2). To visually assess sample viability, cell suspensions were treated with DAPI marker (1:10^4^) prior to image acquisition. DAPI stains the nuclei of dead cells blue through the selective binding of DNA in nonviable cells lacking functional membranes.

### Library preparation and sequencing

Single-cell mRNA sequencing libraries were generated from three biological replicates of age-synchronized wild-type day-one adults and four biological replicates of age-synchronized *daf-2*^-/-^ day-one adults, including two replicates each of the temperature-sensitive *daf-2*^-/-^ *e1370* and *m41* alleles. In addition, one sample of age-synchronized *daf-16*^-/-^; *daf-2*^*-/-*^*(e1370)* day-one adults was sequenced alongside the wild-type and *daf-2*^-/-^ samples.

Prior to loading onto the Chromium™ controller, live cells were quantified with a Bio-Rad TC20™ automatic cell counter. Sample input volumes were calculated following the 10x Genomics^®^ Chromium™ Single Cell 3’ Reagent Kit User Guide (CG00026). Cell viability was assessed using 0.4% trypan blue counterstaining. Each sample was assayed at least twice to increase precision. Our empirical assessments of the reproducibility of live *C. elegans* cell quantifications obtained using the Bio-Rad TC20™ demonstrated that accurate quantification of heterogeneous mixtures of dissociated *C. elegans* cells without the use of FACS is inherently challenging. Cell count measurements obtained with conventional benchtop automatic cell counters not designed for assaying live *C. elegans* cells (eg. Bio-Rad TC20™ and Invitrogen Countess II™) were empirically found to be about an order of magnitude lower than the cell counts obtained using FACS, the actual sc-RNA-Seq results, and calculated predictions based on the number of input worms per sample. Serendipitously, the low input cell concentrations systematically reported by the Bio-Rad TC20™ automatic cell counter resulted in higher-than-expected cell counts in the final sequencing data. The majority of libraries generated contained ∼10-40K individual cells per sample (Figure S3), which is well above the previously reported upper limit of single cells that can be sequenced per sample (https://kb.10xgenomics.com/hc/en-us/articles/360001378811-What-is-the-maximum-number-of-cells-that-can-be-profiled-).

The loading concentration of live *C. elegans* single cells ranged from 50-350 cells / µL when measured using a Bio-Rad TC20™ automatic cell counter with 0.04% trypan blue dye (Table S10). Single-cell 3’-mRNA sequencing libraries were prepared using the 10x Genomics^®^ GemCode™v1.0 Single-Cell 3′ Gel Bead and Library Kit, as described previously (Zheng, Terry et al. 2017). Briefly, cells were loaded onto a 10x Genomics^®^ Chromium™ microfluidics controller at a limiting dilution and encapsulated by water-in-oil emulsion nanodroplets containing Gel beads in EMulsion (GEMs) and reagents. Each GEM contains oligonucleotides which facilitate reverse transcription (RT) of polyadenylated mRNA transcripts and the attachment of the following barcodes and sequencing adapters: (i) 30 bp anchored oligo-dT primers to hybridize with polyA, (ii) 14 bp GEM-specific oligos for single-cell indexing, (iii) 10 bp unique molecular identifier (UMI) oligos for single-molecule indexing, (iv) template-switching oligos which attach to 3’-cDNA during RT to facilitate attachment of Illumina adapter sequences.

After encapsulation, cells were chemically lysed and mRNA was captured via polyA-hybridization to oligo-dT primers to generate single-cell GEMs. This step excludes bacterial RNA contamination via selection of eukaryotic polyA-containing transcripts. GEM reverse transcription (GEM-RT) was performed in a thermal cycler: 55°C for 2 h; 85°C for 5 min; held at 4°C overnight. After GEM-RT, the oil emulsions were broken to combine the barcoded single-stranded cDNAs; the pooled transcripts were cleaned with Mag-Bind^®^ TotalPure NGS Beads (Omega Bio-tek). Purified cDNA was amplified in a thermal cycler with primers complementary to the template-switching oligos and the Illumina sequencing adapters: 98°C for 3 min; 14 cycles x (98°C for 15 s, 67°C for 20 s, and 72°C for 1 min); 72°C for 1 min; held at 10°C. PCR-amplified cDNA was cleaned with Mag-Bind^®^ TotalPure NGS Beads and sheared to ∼200 bp using a Covaris M220 Focused-Ultrasonicator.

Sample indices and adapters were attached using the 10x Genomics^®^ GemCode™v1.0 kit reagents for end-repair, A-tailing, adapter ligation, post-ligation cleanup, sample indexing, and PCR cleanup. The final sequencing libraries were assayed with quantitative PCR (KAPA Biosystems Quantification Kit for Illumina Libraries), and fragment size distributions were measured using an Advanced Analytical Fragment Analyzer (Figure S4). Sequencing libraries were pooled and diluted to a concentration of 2 nM with Qiagen EB buffer (10 mM Tris pH 8.5 with 0.1% Tween 20). Sequencing was conducted on five lanes of an Illumina NextSeq 500 in high-output mode (V2 chemistry, 2 x 75-cycle kits). Single-cell libraries generated using the 10x Genomics^®^ GemCode™v1.0 platform require paired-end sequencing with dual indexing (10x Genomics^®^ Chromium™ Single Cell 3’ Reference Card RevA CG00037). The following read lengths were required to capture the cell, sample, and UMI barcodes from the v1.0 - generated 10x Genomics^®^ GemCode™libraries: 98 bp Read 1, 10 bp Read 2, 8 bp I5 Index, and 14 bp I7 Index. Read 1 captured the cDNA; Read 2 captured the UMIs; Index I5 captured the sample indices; Index I7 captured the GEM (cell) barcodes.

### Sequence alignments and barcode processing

Sequencing reads were demultiplexed and converted to fastq format using the 10x Genomics^®^ Cell Ranger™(v1.2.1) software (https://support.10xgenomics.com/single-cell-gene-expression/software/overview/welcome), which compares cell barcodes against a whitelist and filters mismatches based on quality scores. Reads with identical UMIs and GEM barcodes were identified as PCR duplicates and removed. Transcriptomic cDNA sequences were aligned to the *C. elegans* reference genome PRJNA13758 (WS256) using the RNA-Seq aligner STAR using default settings (Dobin, Davis et al. 2013) implemented via the Cell Ranger™(v1.2.1) software pipeline. Gene annotations were filtered to include only protein-coding genes. Sequencing reads that mapped confidently to exonic regions of protein-coding genes and were correctly associated with valid sample, cell, and molecular barcodes were included in the final gene-cell-barcode matrix. Data generated from multiple sequencing runs of a single library were merged into one dataset using Cell Ranger™(v1.2.1); duplicate reads were excluded.

Normalized digital gene expression matrices were generated for each sample using Seurat (v1.4.0), as described previously (Satija, Farrell et al. 2015). Briefly, the filtered gene-cell-barcode matrices generated with Cell Ranger™(v1.2.1) were normalized to a total of 10^4^ UMIs and log-transformed using Seurat (v1.4.0). Cells with fewer than 50 genes or greater than 5% mitochondrial sequences were excluded from the analysis. UMIs present in fewer than 3 cells were excluded, and sequencing errors were reduced by eliminating mismatched bases in reads with identical UMIs. Cells with more than 1,500 UMIs were removed after visual inspection of gene count distributions in order to eliminate potential cell doublets (Figure S5). To improve downstream cell clustering (Buettner, Natarajan et al. 2015), cell-to-cell variation in the gene expression data due to batch effects was regressed out using Seurat-implemented negative-binomial models, with the number of molecules detected per run set as the confounding variable. Gene expression matrices were generated from each library individually and saved as Seurat (v1.4.0) objects using R (v3.4.1).

To verify that the sequencing reads mapped appropriately to the 3’-ends of *C. elegans* protein-coding mRNA transcripts, bam files were indexed with Samtools and visualized using the Integrative Genomics Viewer (Robinson, Thorvaldsdottir et al. 2011, Thorvaldsdottir, Robinson et al. 2013) (Figure S6).

### Cell-correlated sample alignment

To perform an integrated analysis of cell-specific differential gene expression based on *daf-2* status, an unsupervised, machine-learning-based dataset alignment of 7 independent samples containing a total of 115,000 wild-type and *daf-2*^*-/-*^ cells was performed using Seurat (v2.0.1) Canonical Correlation Analysis (CCA), as described in (https://satijalab.org/seurat/Seurat_AlignmentTutorial.html). The Seurat (v2.0.1) CCA implements algorithms to identify and align common cell types based on the co-correlation of gene expression matrices across multiple datasets (Butler, Hoffman et al. 2018), allowing directly automated comparisons of gene expression profiles for specific cell types across genotypes. Prior to sample alignment, datasets were reformatted in R (v3.4.2) as follows: (i) Seurat (v1.4.0) objects were updated to allow compatibility with Seurat (v2.0.1) software, (ii) sample IDs were appended to each sample’s cell barcodes to avoid downstream data processing errors caused by duplicate cell barcodes, and (iii) metadata columns were appended to the expression matrices recording the sample ID, library ID, *daf-2*^*-/-*^-status, UMI count, strain name, genotype, and sequencing date of each cell to allow downstream tracking of cell parameters after the datasets were merged.

Canonical correlation vectors (CCs) describing the expression covariance across datasets were calculated using Seurat (v2.0.1) RunCCA with default parameters; the 570 most variable genes were included in the CC analysis. The statistical significance of the CCs was assessed graphically using the Seurat (v2.0.1) PCHeatmap and PCPlot functions (Figure S7), as well as the Seurat (v2.3.4) MetageneBicorPlot (MBP) function (Figure S8). The MBP function plots the correlation strength of each CC in order to determine the number of CCs required to saturate the linear relationship between CC quantity and correlation strength of across genotypes. The top 120 CCs demonstrated a significant contribution to genotype correlation strength based on the existence of a downward slope in the MBP plot around CC 120. The Seurat (v2.0.1) RunCCA dataset alignment procedure was executed numerous times using a range of alternative values of CCs to assess the relationship between CC quantity and cluster identities, and qualitatively similar results were obtained for CCs ∼100-140.

The top 120 CCs calculated by Seurat (v2.0.1) were used to project each individual dataset into a maximally-correlated subspace and to then combine the datasets in a dimensionally-reduced subspace grouped into local communities based on similar gene expression patterns. Cells were excluded from the downstream analysis if the variance explained by the CCA was less than half that explained by a standard Principal Component Analysis (PCA) (Butler, Hoffman et al. 2018). The final merged dataset contained ∼40,000 cells computationally partitioned into 101 mixed-genotype cell-type-specific clusters. The pooled, aligned, and annotated samples were saved as a single Seurat (v2.0.1) object, enabling a high-throughput and integrated analysis of every single-cell transcriptome across all samples

### Cell clustering analysis

Principal component analysis (PCA) was performed on the pooled dataset containing ∼40,000 aligned wild-type and *daf-2*^*-/-*^ cells using Seurat (v2.0.1) to calculate the multidimensional vectors defining gene expression co-variance patterns across all cells(Satija, Farrell et al. 2015). Briefly, genes with highly-variable expression were selected by Seurat based on dispersion (expression variance / mean) z-scores (Macosko, Basu et al. 2015), which take into account the relationship between gene expression and variability. The genes with the most variable expression across all single cells were detected using Seurat (v2.0.1) FindVariableGenes with default parameters. The top 2100 genes were used to calculate the principal components (PCs) from the normalized gene expression matrices. The significance of the calculated gene expression variance parameters was validated based on visual inspection of the dispersion plot (Figure S9).

Seurat RunPCA detected a total of 568 PCs describing the gene expression co-variation across all cells in the dataset. The statistical significance of the top PCs was assessed graphically using the Seurat (v2.0.1) PCHeatmap, PCPlot, and PCElbow functions (Figure S10, S11). The PCElbow plot enables visualization of the standard deviation (SD) associated with each PC in order to identify the minimum number of principle components required to saturate the relationship between variance and PC quantity. This approach is known conventionally as the so-called “elbow” graphical method, a commonly used technique for assessing the statistical significance of cluster parameters (Zhang, Lee et al. 2018).

Cell clustering was performed with Seurat (v2.0.1), which uses graphical-embedding algorithms to partition cells into local communities based on gene expression similarity, as described previously (Waltman and van Eck 2013, Macosko, Basu et al. 2015). Briefly, cells were computationally embedded in a shared nearest neighbor (SNN) graphical structure based on their Euclidean distance in PC-space. Edges were drawn between similar cells with edge weights calculated based on local overlap (Jaccard distance) (Levine, Simonds et al. 2015). The Louvain modularity optimization algorithm was implemented to iteratively partition cells into communities based on edge weights (Blondel, Guillaume et al. 2008). The Louvain algorithm uses the local moving heuristic approach to randomly analyze the nodes (cells) in a community (tissue) in a random order and to move the nodes repeatedly until maximum modularity is achieved in the dataset. The Jaccard index stringency cutoff used for edge pruning during SNN construction was set at 0.0667 and the k-nearest neighbor parameter was defined as 30. The number of random starts used to optimize cluster modularity was 100, with a maximum of 10 iterations used per random start. The cell clustering procedure was repeated multiple times using a range of alternative PC dimensions and qualitatively similar results were obtained using PCs ∼100-140, leading to the selection of the top 120 of the total 568 PCs for determining the final clusters. The resolution parameter of FindClusters was set at 3.0 to optimize cluster granularity.

### Identification of cluster-specific biomarkers and functions

Biomarker genes defining each cell cluster were identified using Seurat (v2.0.1) FindMarkers, which determines the unique gene expression profile of each cluster based on differential gene expression comparisons across clusters. Cell type biomarkers common to the wild-type and *daf-2* samples were identified in addition to biomarkers defining only the wild-type and daf-2 cells within each cluster. To be included in the analysis, genes were required to be detected in at least 1% of cells in each sample and to have average differential expression levels greater than 1% between the two samples.

The *C. elegans* gene annotation Bioconductor package org.Ce.eg.db (v3.6.0) was used to convert WormBase (WS256) GeneIDs to standard gene names (Gentleman, Carey et al. 2004). Functional identities were assigned to the machine-generated clusters using the conventional supervised approach, as described in (Villani, Satija et al. 2017). Cell clusters were annotated with anatomical and functional identities based on the expression of canonical tissue-specific *C.elegans* marker genes identified using the Wormbase tissue expression database(Stein, Mangone et al. 2001, Stein, Sternberg et al. 2001, Chen, Harris et al. 2005, O’Connell 2005).

### Calculating differential tissue-specific gene expression between wild-type and *daf-2* worms

Unbiased machine-learning based identification and alignment of common cell types across samples into a single merged dataset allows for an integrated quantitative analysis of cell-specific transcriptional responses to organism-wide temperature-sensitive knockdown of *daf-2*. During the cell-correlated sample alignment pipeline (described above), gene expression measurements for each cell were globally scaled and normalized across the age-synchronized wild-type and *daf-2*^*-/-*^ single-cell transcriptomes using the Seurat (v2.0.1) LogNormalize function, which normalizes gene expression by total expression, multiplies by a scale factor of 10,000, and then log-transforms the result. After scaling, log-normalization, and alignment of gene-expression data across all samples and cell types, statistically significant cell-specific differential gene expression patterns can be directly calculated using a single, integrated high-throughput pipeline.

Systems-level quantification of statistically significant differential gene expression based on genotype was performed on the 101 aligned, mixed-genotype cluster was performed using the Seurat (v2.0.1) FindMarkers function, which calculates differential expression between any two groups of cells using the non-parametric Wilcoxon rank sum test. Genes with significantly altered expression between wild-type and *daf-2*^*-/-*^ transcriptomes were calculated for each cluster individually using a threshold of padj < 0.10 (∼ p << 0.001) to define significance. Biomarkers were determined for wild-type and daf-2 cell subsets independently as well as combined (Supplemental File 3). The unique transcriptional response of each cluster due to the knockdown of *daf-2* was compared with all other clusters to differentiate between the tissue-specific and global effects of *daf-2* knockdown.

### Non-linear dimensional reduction for t-SNE visualization

Single-cell clusters were visualized with Seurat (v2.0.1) using t-Distributed Stochastic Neighbor Embedding (t-SNE), a method of non-linear dimensionality reduction for visualizing high-dimensional cell relationships in low-dimensional space (van der Maaten and Hinton 2008, van der Maaten 2014). The significant PCs identified during clustering analysis were used as input to generate the t-SNE plot, which is a two-dimensional embedding of the multi-dimensional data. In a t-SNE plot each cell is represented by a dot, and the proximity of the dots correlates with the transcriptional similarity of the cells. Marker gene expression was plotted in the t-SNE plots using normalized UMI counts.

### Experimental Design and Rationale

It is exceedingly important to demonstrate the logical coherency of a single-cell datasets prior to making any assessments of tissue-specific gene expression dynamics. Meaningful interpretations of any transcriptomic dataset are only made possible through the persistent incorporation of a variety of approaches to meticulously assess data significance and logic throughout the entire protocol. In order to confirm and support the congruency of the dataset. numerous quality control tests and sanity checks for logical consistency were performed throughout the analysis.

To assess data reproducibility across time, library preparation and sequencing were performed in two separate rounds, separated by four months’ time. Sequencing libraries were generated for at least two biological replicates per genotype using unique starting populations of worms. Four libraries were prepared and sequenced simultaneously during the first round, and five libraries were prepared and sequenced simultaneously during the second round. Biological replicates from each genotype were evenly distributed across runs to reduce sample prep bias. To ensure that biological replicates were well-distributed across the aligned sample-specific clusters, t-SNE plots were generated with the cell dots colored according to their library of origin (Figure S12).

To visually confirm sample correlation globally across biological replicates, whole-organism gene expression heatmaps and dimensionality reduction plots were generated using the Comprehensive R Archive Network (CRAN) package gplots (v3.0.1) (Figure S12). Gene counts of protein-coding genes were calculated from indexed bam files using HTSeq-count and then differential gene expression was quantified using the DESeq2 package from Bioconductor (Gentleman, Carey et al. 2004, Love, Huber et al. 2014). IGV plots were generated to visually confirm upregulation of Class I longevity genes in the *daf-2*^-/-^ samples (Figure S13).

As an additional sanity check on differential expression measurements, *C. elegans* tissue-specific biomarker profiles and transcriptional and responses to global *daf-2* knockout were calculated directly from the raw sequencing data using a supervised approach. Cell-specific expression results were normalized by the number of molecules using UMI tags and compared across samples using a supervised approach. Tissue-specific differential expression was visualized with volcano plots (Figure S14).

### Raw Datasets

The final dataset is available for download as a Seurat R object:

https://pages.uoregon.edu/jpreston/WTall_DAF2all_names_numbers_120_R3_2018.Rda.

To request raw data files and scripts, please.

## Supplemental Data

**Supplemental File 1.** Tables S1-S10.

**Table S1**. **Cluster Biomarker Signatures.** Each machine-generated cell type cluster is characterized by the expression of a unique group of biomarkers which is stable across replicates.

**Table S2. Cluster Cell Counts by Genotype.** The cell type clusters in the aligned dataset are comprised of a heterogeneous mixture of cells from seven independent populations of age-synchronized wild-type and *daf-2*^*-/-*^ worms.

**Table S3. Single-Genotype Clusters**. Twenty of the smaller clusters contained cells of a single genotype rather than a mixture of wildtype and daf-2 cells.

**Table S4. Predicted Cell Types**. Functional identities were assigned to each cluster using a supervised approach based on matching the machine-generated biomarkers to known canonical *C. elegans* tissue-specific genes reported in the literature.

**Table S5. *C. elegans* Biomarker Genes.** Cell-specific biomarkers annotated with references and documented expression (partial list).

**Table S6. Functional Gene Networks Involved in Tissue-Specific *Daf-2*^*-/-*^ Loss.** Significantly altered expression was detected in genes involved in several age-related physiological processes.

**Table S7. Global *Daf-2*-Response Genes.** Twenty-seven genes were identified that are significantly differentially expressed in every cell type.

**Table S8. Transcriptional Response of Epidermal Seam Cells to *Daf-2*^*-/-*^.** Seam cells displayed the strongest response to the loss of *daf-2* in terms of the number of significantly upregulated age-related genes.

**Table S9. Sperm-Specific Genes with Altered Expression in *Daf-2*^*-/-*^ Tissues.** Many germline and sperm-specific genes were upregulated in somatic tissues of *daf-2*^*-/-*^ worms.

**Table S10. Library Preparation, Alignment, and Cell Count Statistics.** The loading concentration of live *C. elegans* single cells ranged from 50-350 cells/µL when measured using a Bio-Rad TC20™ automatic cell counter. Sequencing reads that mapped confidently to exonic regions of protein-coding genes and were correctly associated with valid sample, cell, and molecular barcodes were included in the final gene-cell-barcode matrix.

**Supplemental File 2.** Genes with altered expression between wild-type and *daf-2*^*-/-*^ by cluster.

**Supplemental File 3.** Genotype-specific biomarker genes.

**Supplemental File 4.** Figures S1-S14.

### Acknowledgements

This work was made possible by Mary and Tim Boyle’s 2016 donation in support of basic genomics research at the University of Oregon. The 10X Genomics Chromium™instrument and the associated consumables that were used to prepare the samples described in this work were purchased with a portion of the gift funds. Sequencing runs and reagents were funded by NIH Grant AG056436. We thank Doug Turnbull, John Willis, Jason Sydes, Peter Batzel, Jim Stapleton, Anna Coleman-Hulbert, and Ash Wilson for comments. We thank WormBase.

## Author Contributions

Conceived and designed the experiments: JLP. Performed the experiments: JLP, MW. Analyzed the data: JLP, NS. Contributed reagents/materials/analysis tools: NS, MW. Wrote the paper: JLP. The authors declare no competing interests. All authors have read and approved this manuscript. This article is the authors’ original work and has not been submitted for publication elsewhere. JLP takes full responsibility for the validity and legitimacy of the data and its interpretation.

