## Supplemental File 4 for "Organism-wide single-cell transcriptomics of long-lived *C. elegans daf*-2^-/-^ mutants reveals tissue-specific reprogramming of gene expression networks"

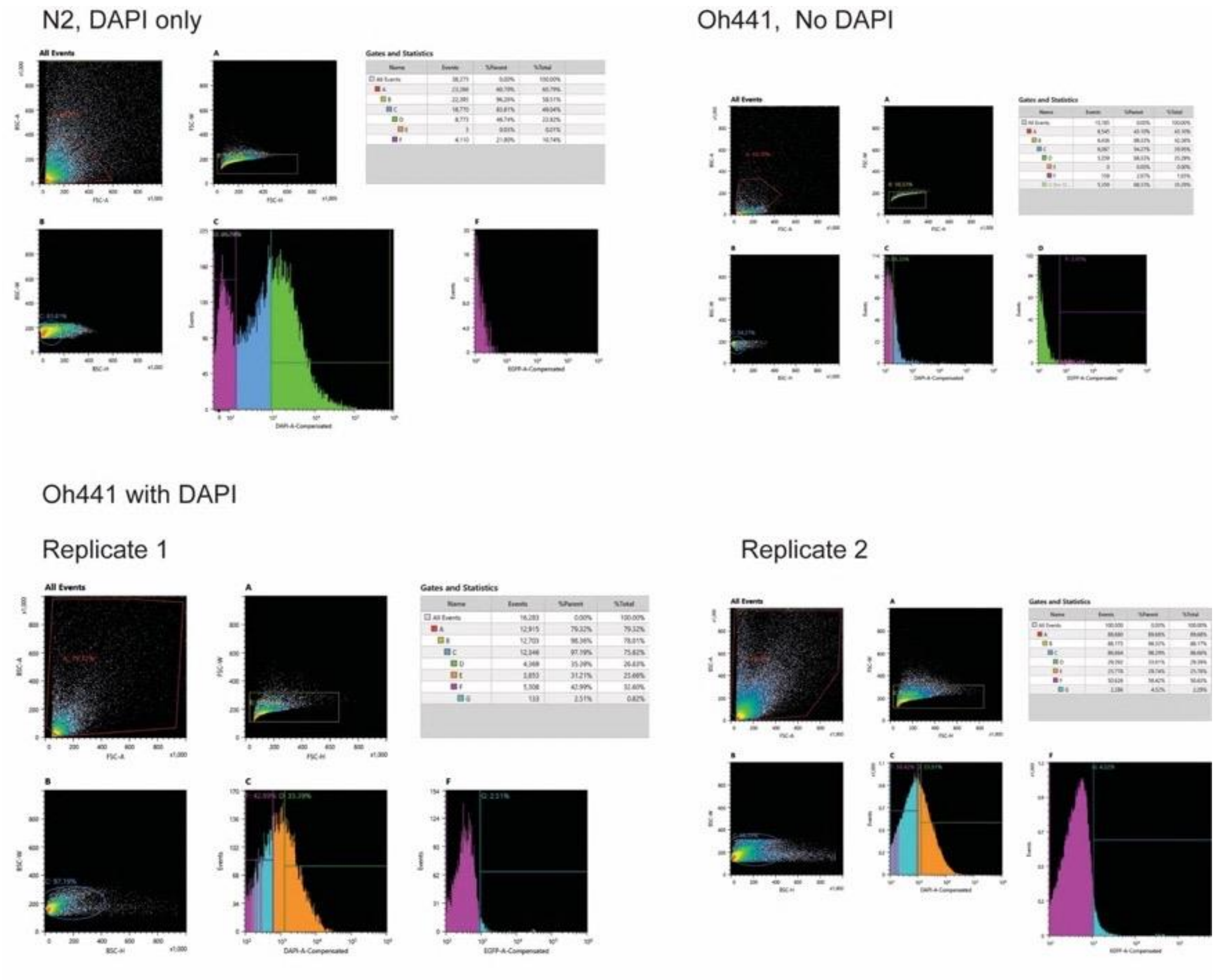


**Figure S1.** FACS gates and collection statistics. Fluorescence-Activated Cell Sorting (FACS) was performed on dissociated cells from adult *C. elegans* expressing GFP under the control of the *unc-119* promoter (OH441: otIs45 [P*unc-119::GFP*]). Cells expressing *unc-119* mRNA were isolated based on high GFP expression ($GFP^{high}$). Cell viability was assessed using DAPI (4',6-diamidino-2-phenylindole, $1{:}10^4$).

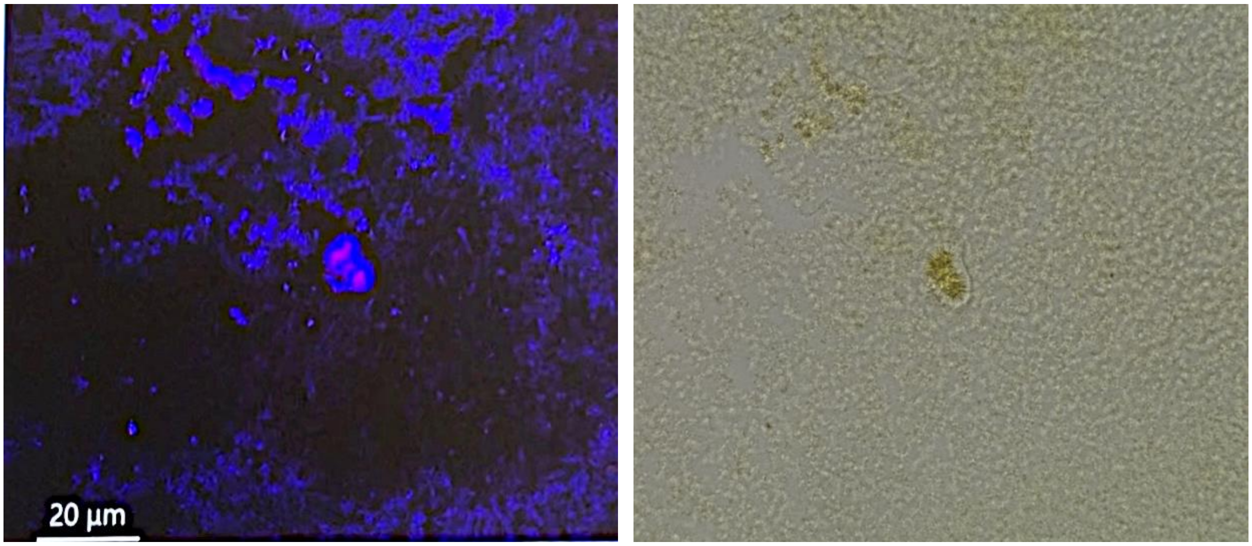


**Figure S2.** Filtered cell suspensions were observed using a Keyence BZ-X800 inverted fluorescent microscope. Cell viability was assessed visually using DAPI (1:$10^4$), which marks nuclear DNA of dead and broken cells with a blue stain.. Photo credit: Derek Johnson.

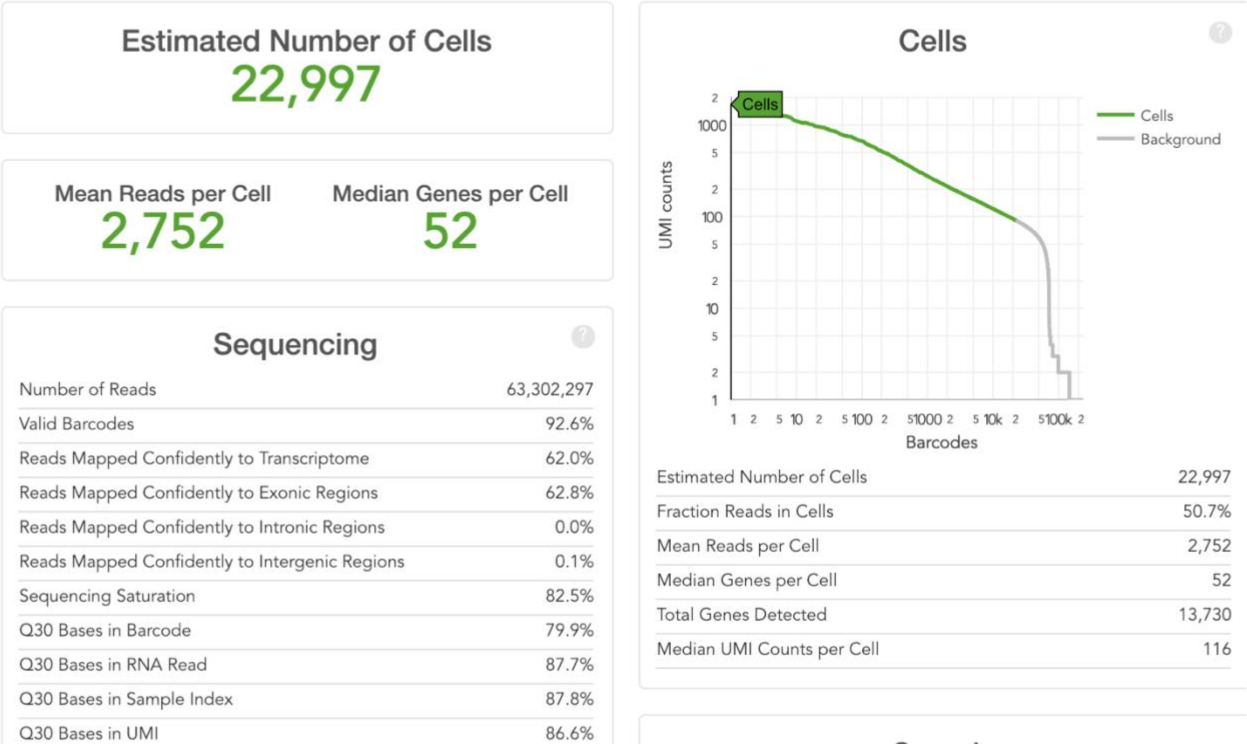


**Figure S3**. Example cell count and barcode alignment statistics. The majority of libraries generated contained ~10-40K individual cells per sample

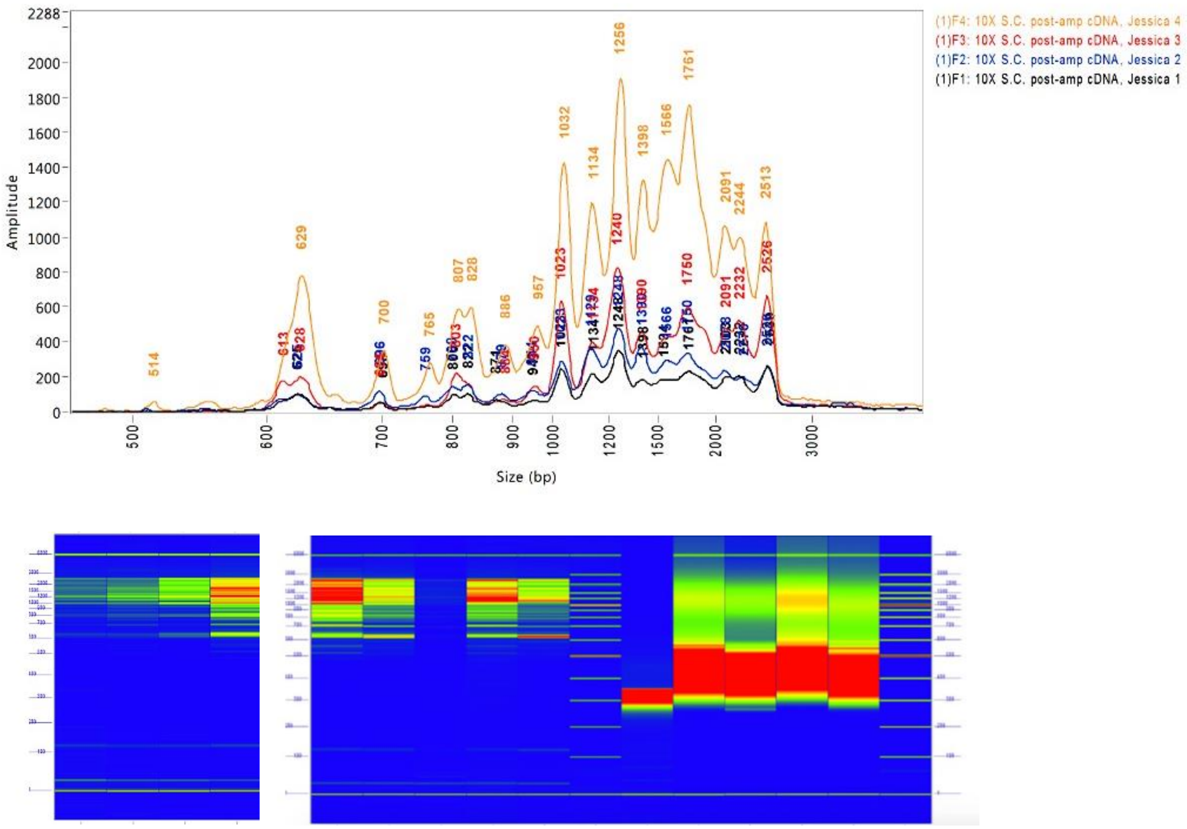


**Figure S4**. The final sequencing libraries were assayed with quantitative PCR (KAPA Biosystems Quantification Kit for Illumina Libraries (top image). Fragment size distributions were measured using an Advanced Analytical Fragment Analyzer (bottom image).

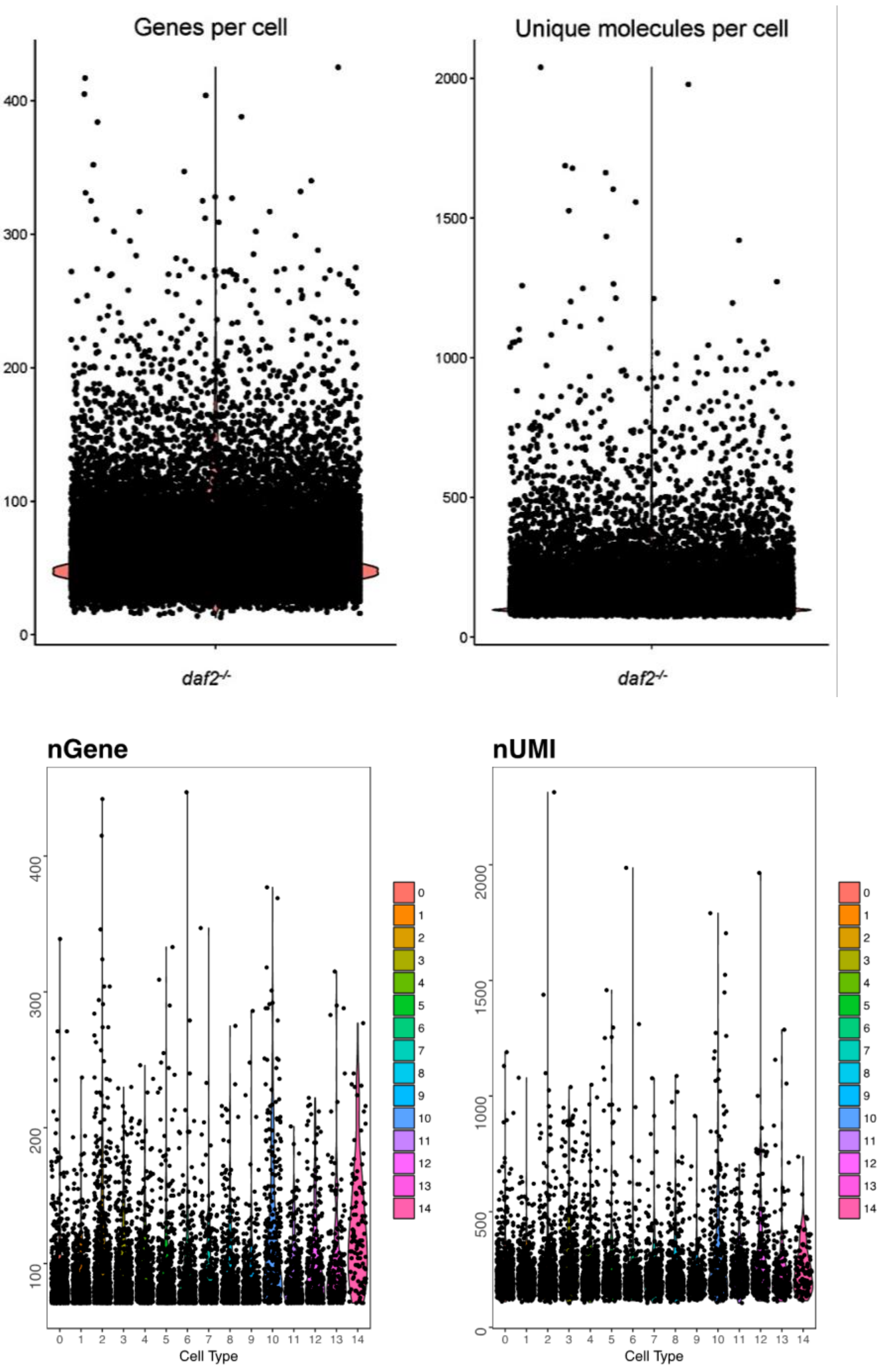


**Figure S5.** Genes and unique molecules per cell. Cells with fewer than 50 genes or greater than 1,500 molecules (UMIs) were excluded from the removed after visual inspection of gene count distributions. Sequencing errors were reduced by eliminating mismatched bases in reads with identical UMIs.

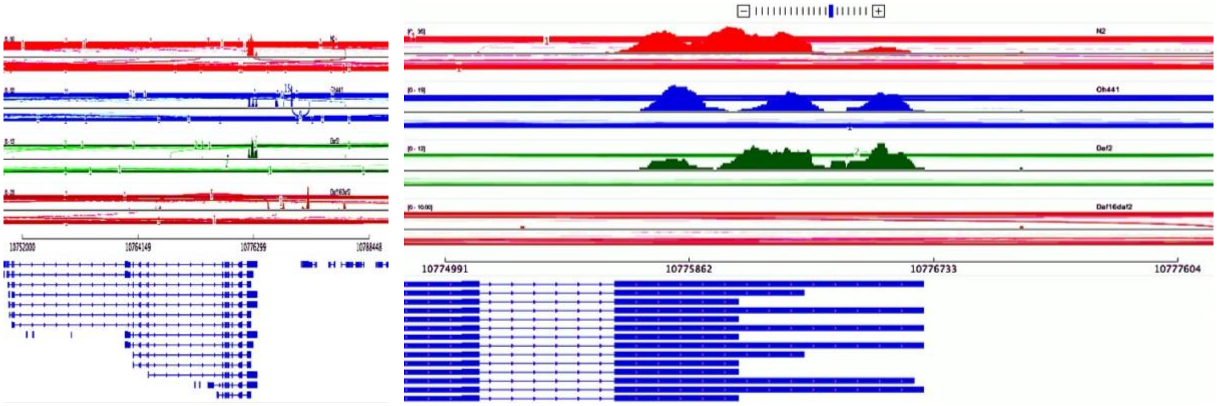


**Figure S6**. To verify that sequencing reads mapped appropriately to the 3'-ends of *C. elegans* protein-coding mRNA transcripts, the *daf-16* locus visualized using sushi plots generated with the Integrative Genomics Viewer (Broad Institute). The bottom track depicts sequencing reads from *daf-16*$^{-/-}$ mutant worms.

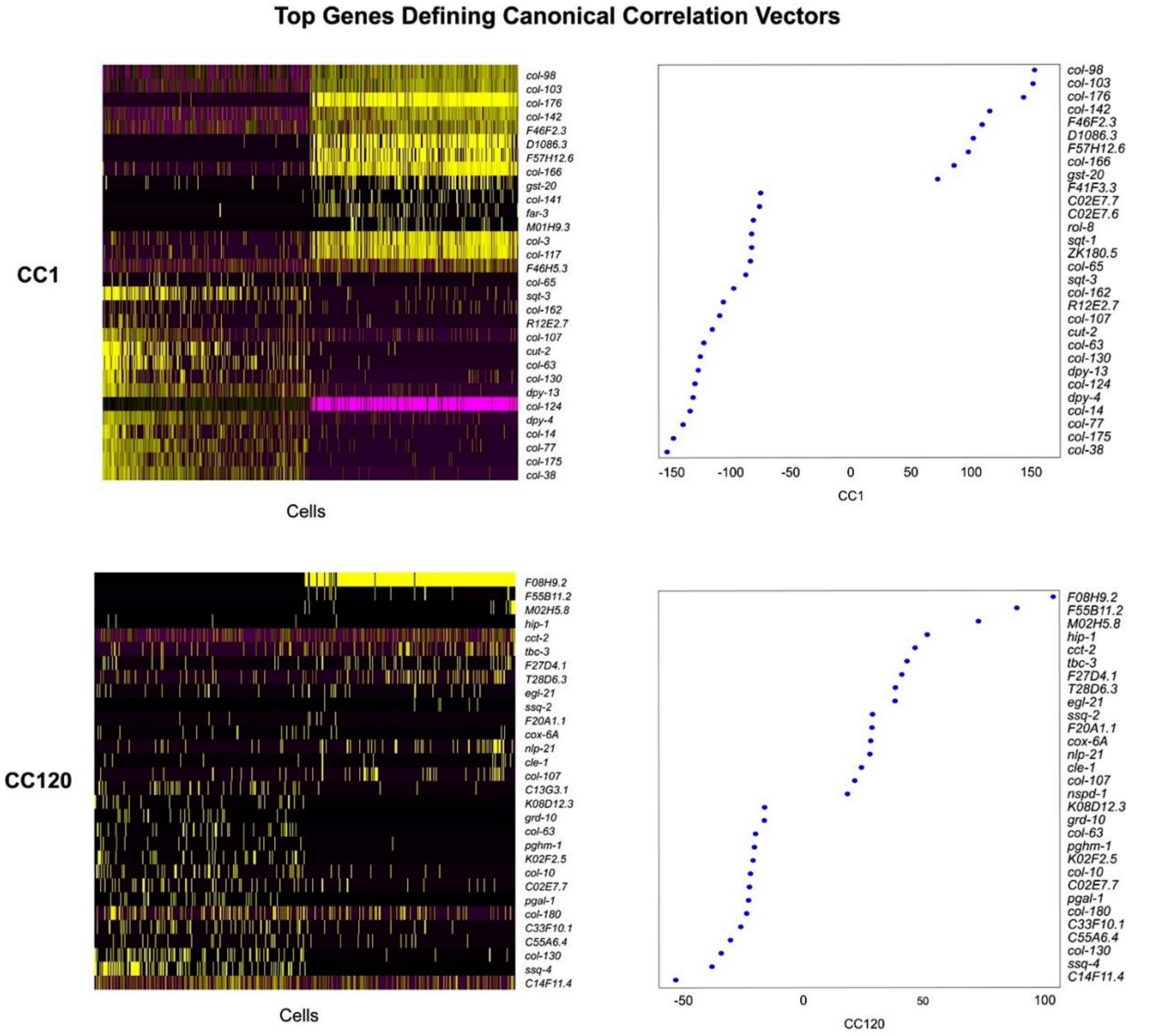


**Figure S7.** Canonical correlation vectors (CCs) describing the expression covariance across datasets were calculated with Seurat (v2.0.1) RunCCA using default parameters. The statistical significance of the CCs was assessed graphically using the Seurat (v2.0.1) PCHeatmap and PCPlot functions.

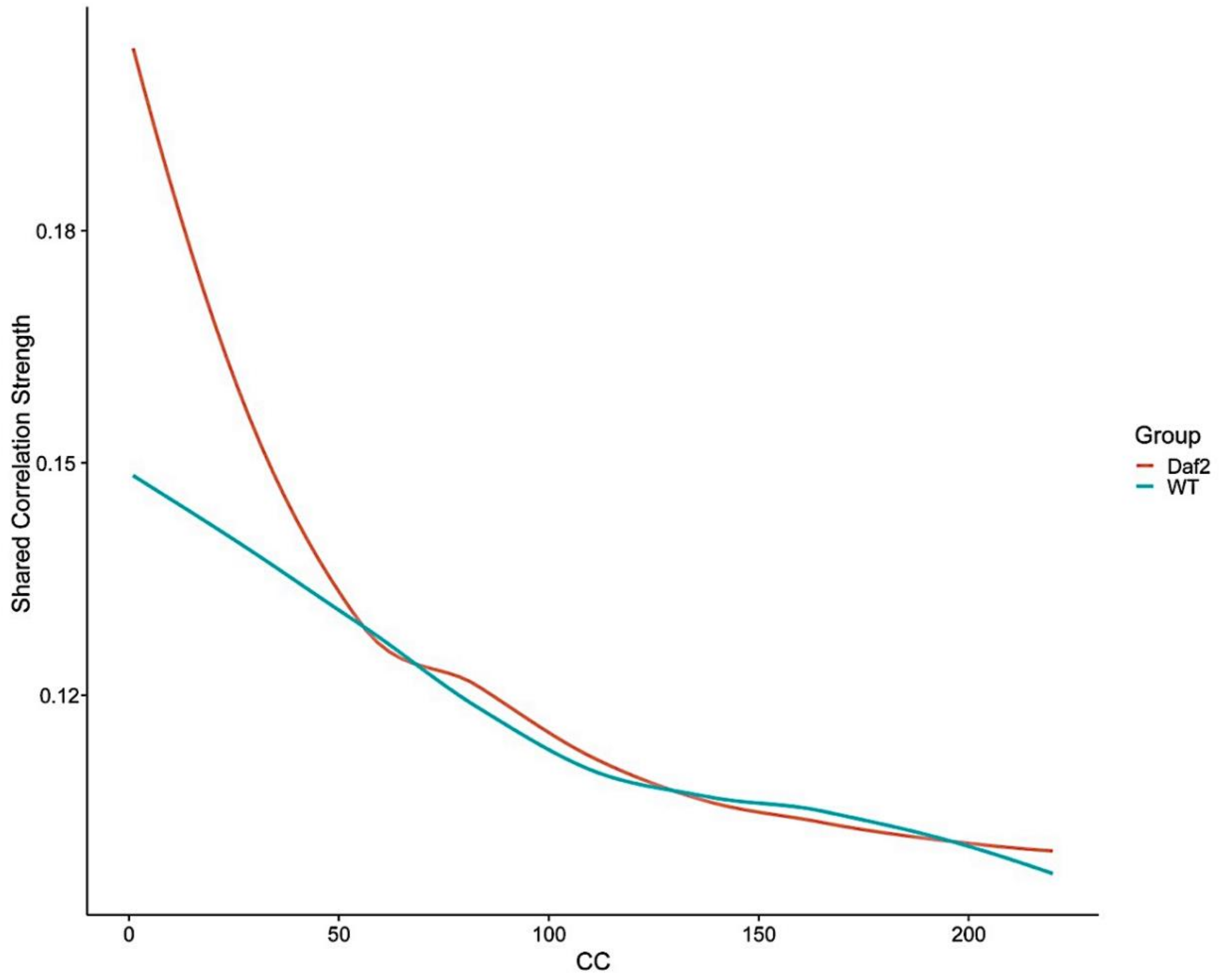


**Figure S8**. The MetageneBicorPlot (MBP) function plots the correlation strength of each CC in order to determine the number of CCs required to saturate the linear relationship between CC quantity and correlation strength of across genotypes. The top 120 CCs demonstrated a significant contribution to genotype correlation strength based on the existence of a downward slope in the MBP plot around CC 120.

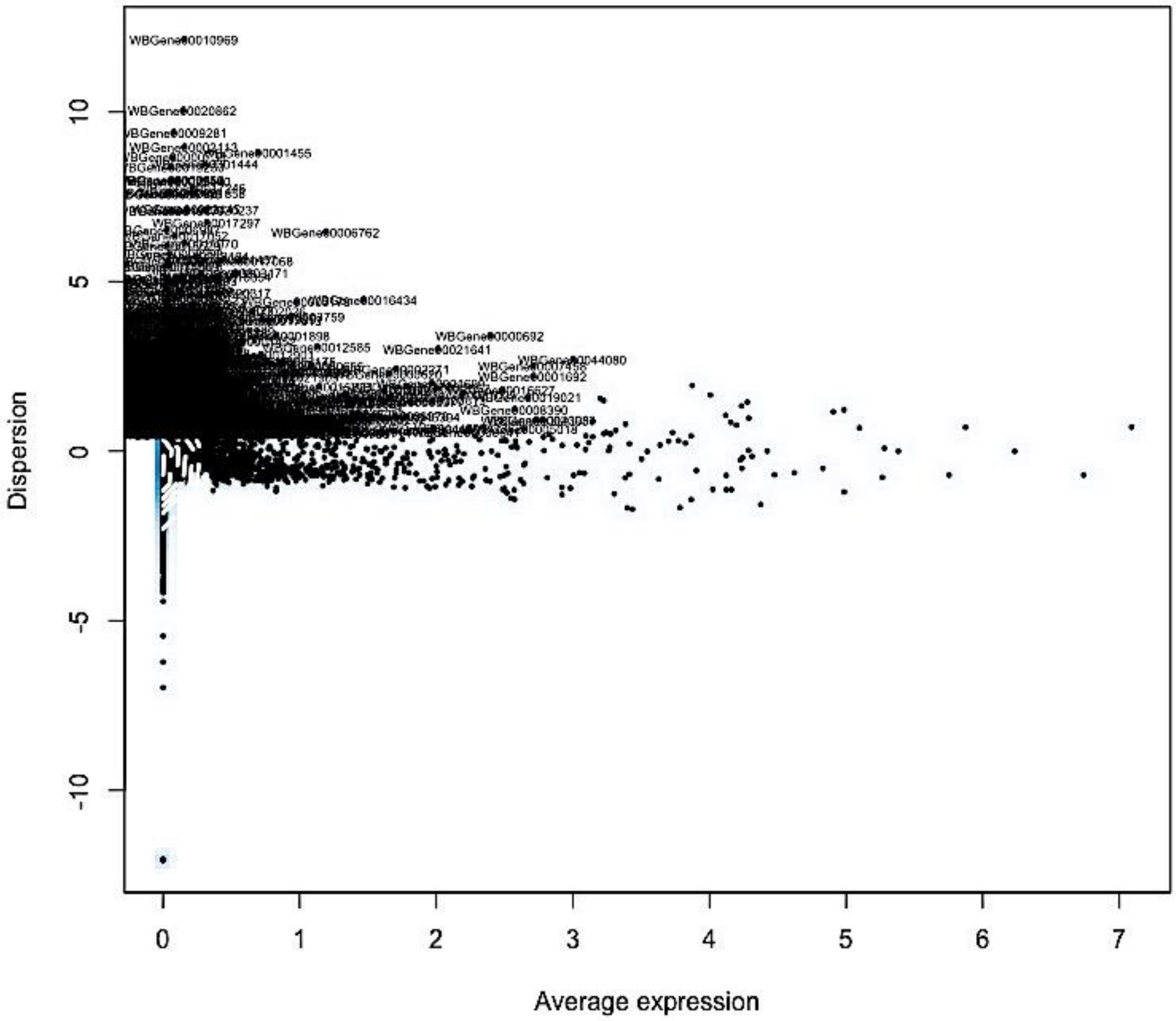


**Figure S9.** Dispersion plot of variable gene expression across all single cells. Genes with highly-variable expression were selected by Seurat (v2.0.1) FindVariableGenes based on dispersion (expression variance / mean) z-scores, which take into account the relationship between gene expression and variability.

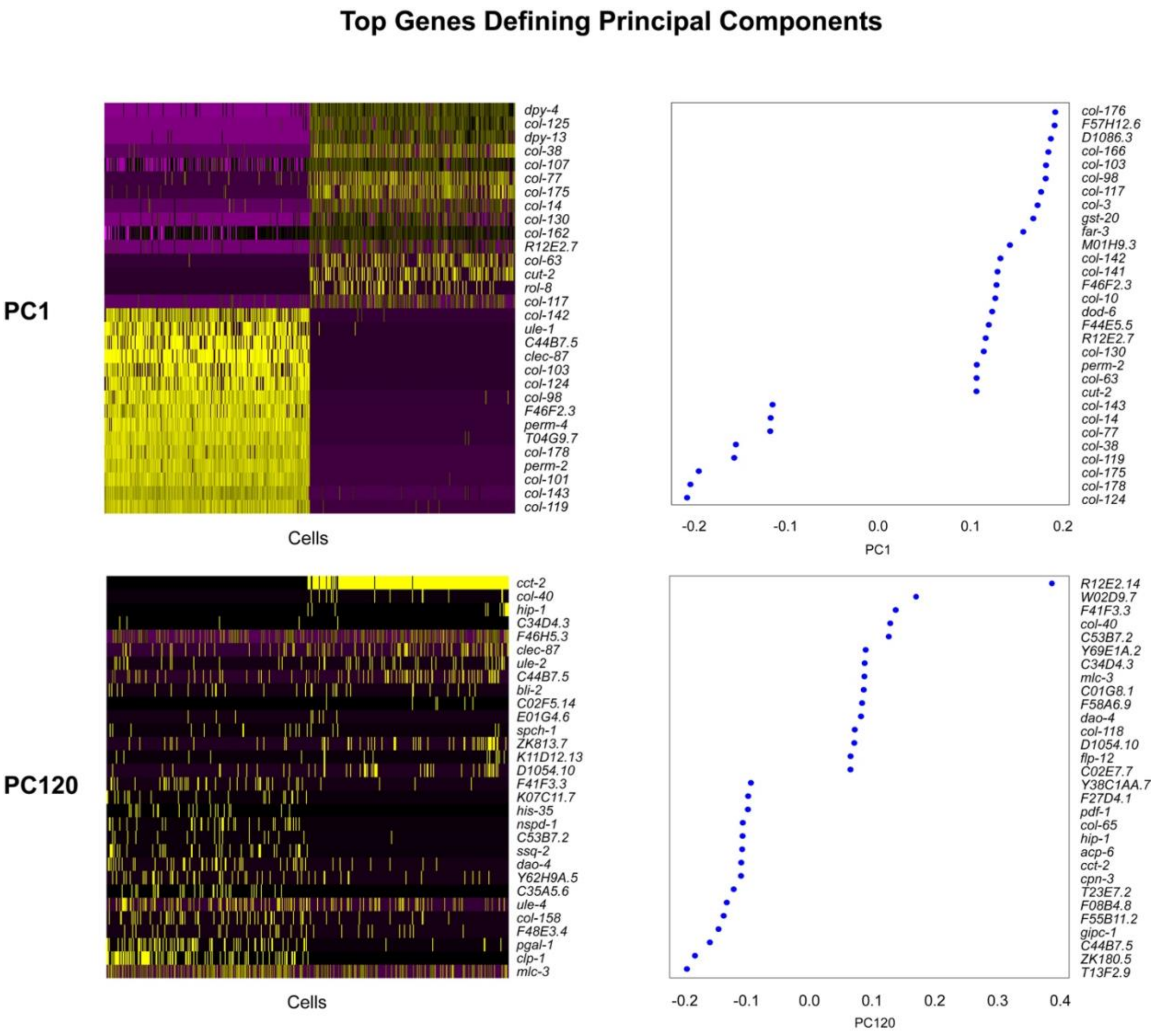


**Figure S10.** Seurat RunPCA detected 568 PCs describing the gene expression co-variation across all cells in the dataset. The statistical significance of the top 120 PCs was assessed graphically using the Seurat (v2.0.1) PCHeatmap and PCPlot functions. The top 120 most significant PCs of the 568 total were included in the downstream cell clustering analysis.

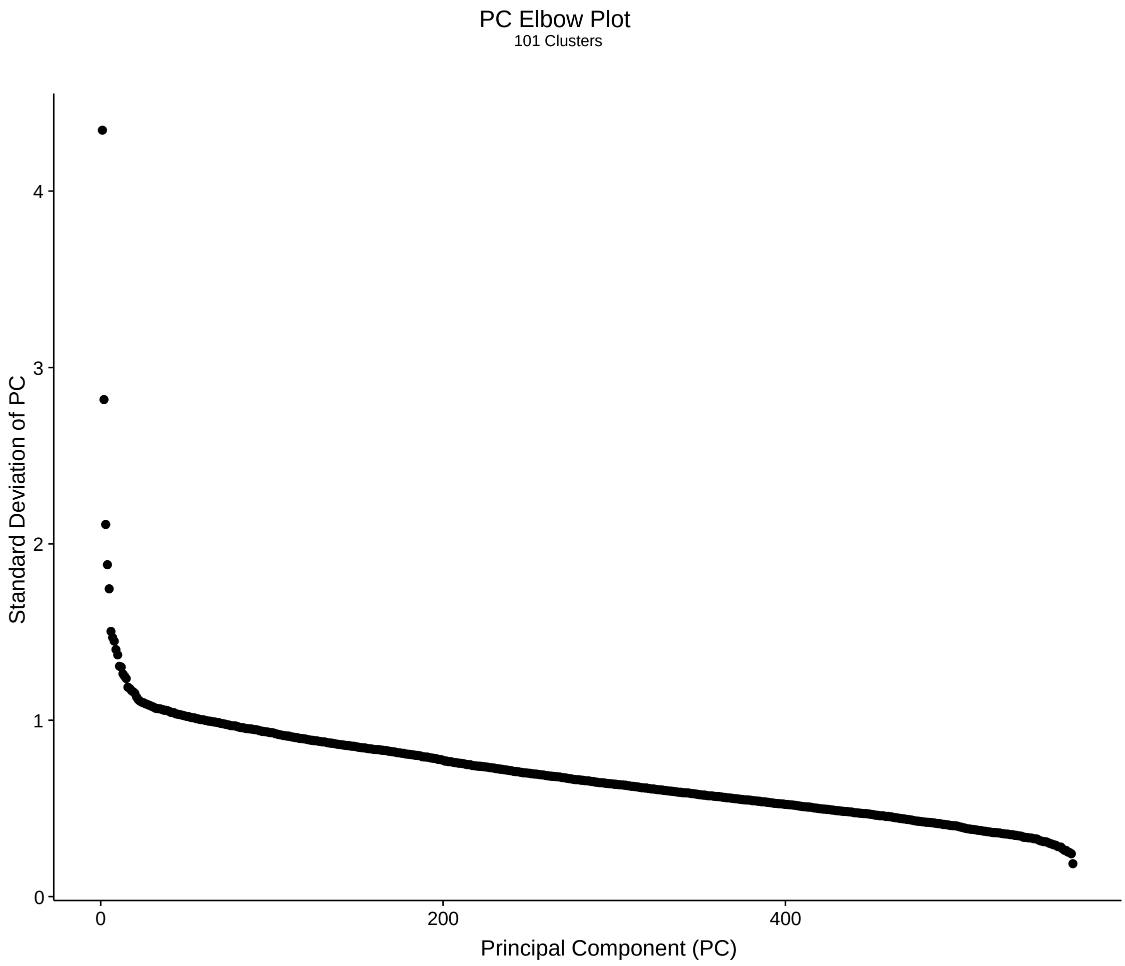


**Figure S11.** The PC Elbow plot enables visualization of the standard deviation (SD) associated with each PC in order to identify the minimum number of principle components required to saturate the relationship between variance and PC quantity. The approach is known as the "elbow" method. Qualitatively similar clustering results were obtained using PCs ~100-140. The top 120 most significant PCs of the 568 total were included in the downstream cell clustering analysis.

**A**. **B.**

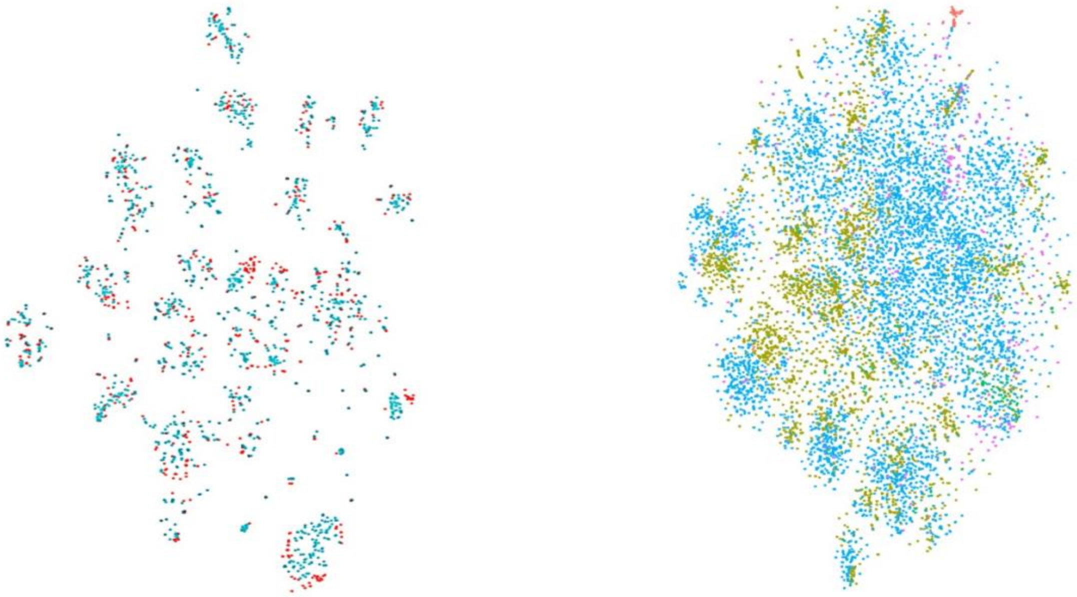

**Figure S12**. t-SNE plots comparing biological replicates to ensure equal distribution of cells from different replicates and sequencing runs within each cluster. **A**. Two replicates of age-synchronized wild-type N2 cells clustered together (pink and cyan dots). **B.** Two replicates of *daf-2*$^{-/-}$(e1370) cells (blue and purple dots) clustered together with two replicates of *daf-2*$^{-/-}$(m41) cells (green and red dots).

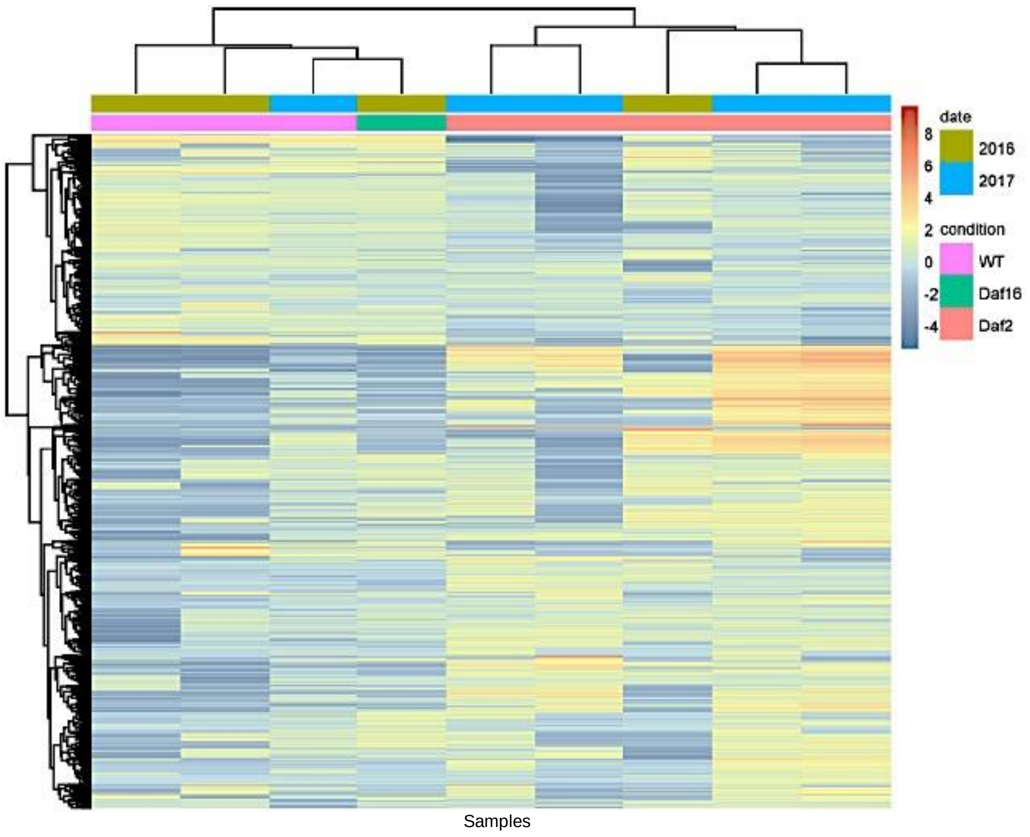


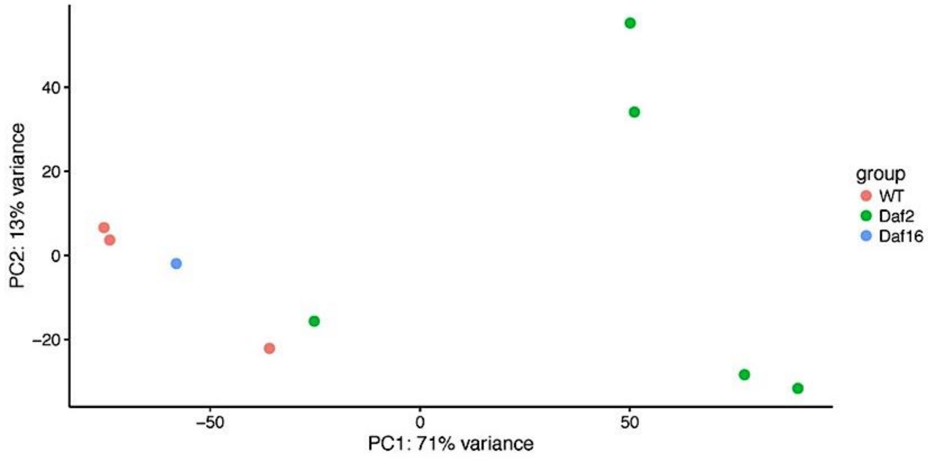


**Figure S13.** Whole-organism gene expression heatmaps and PCA plots were generated to visually confirm sample correlation globally across biological replicates and time. library preparation and sequencing were performed in two separate rounds, separated by four months' time. Sequencing libraries were generated using at least two biological replicates per strain, and replicates from each genotype were evenly distributed across runs to reduce sample prep.

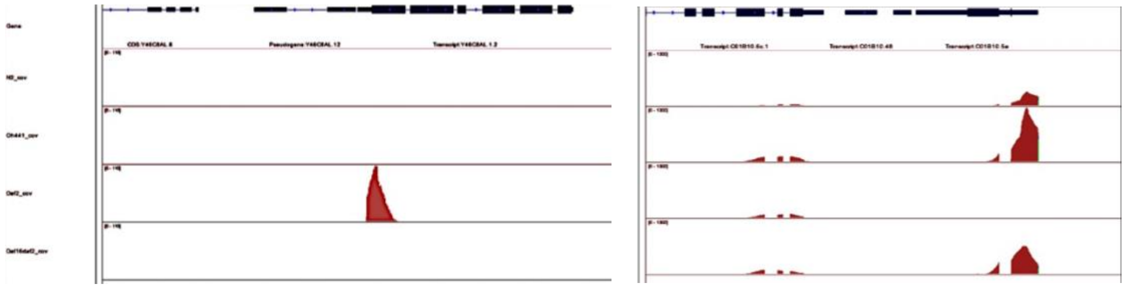

**Figure S14.** To verify that the sequencing reads mapped appropriately to the 3'-ends of *C. elegans* protein-coding mRNA transcripts, bam files produced by Cell Ranger (v1.2.0) were indexed with Samtools and visualized using the Integrative Genomics Viewer (Broad Institute.) IGV plots were generated to confirm upregulation of Class I longevity genes in the daf-$2^{-/-}$ samples

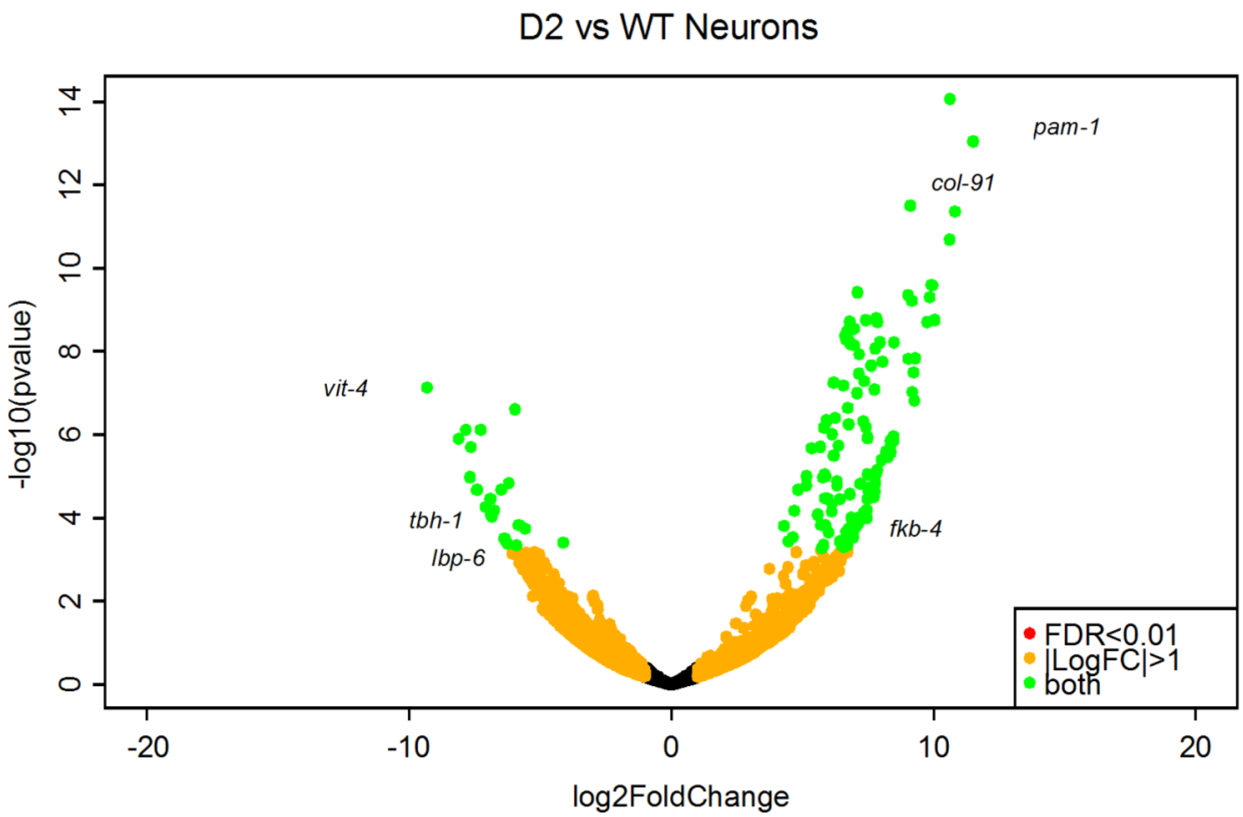


**Figure S15.** As an additional confirmation of internal data consistency, *C. elegans* tissue-specific biomarker profiles and transcriptional and responses to global *daf-2* knockout were calculated directly from the raw sequencing data.
